# Neutrophil lectin-like oxidized low-density lipoprotein receptor (LOX-1) limits inflammatory injury during bacterial pneumonia

**DOI:** 10.64898/2026.11.25.767411

**Authors:** F.T. Korkmaz, Q. Li, E.S. Scholl, M. Mahjabin, D.C. Gatica, R.S. Illig, N. Estrada-Hernandez, F. Moreira-Santana, S.N. Fett., A.G. Hake, L.S. Frye, M.R. Kafka, M. Perez-Vasquez., T. Sawamura, J. Mehta, N.C. Crossland, A. O’Connell, A. Tseng, C.S. Forconi, T. Beach, K.T. Traber, J.P. Mizgerd, L.J. Quinton

## Abstract

Pneumonia is an inflammatory condition that is caused by myriad microorganisms, necessitating broadly applicable therapeutics. Yet, host determinants of pneumonia severity remain poorly understood, demanding a more complete understanding of the cells and signals that control tissue homeostasis. Our group previously showed that the class E scavenger receptor lectin-like oxidized low-density lipoprotein receptor-1 (LOX-1) limits pneumonia-induced lung injury without affecting bacterial clearance, contrasting its well-established injurious effects in the setting of atherosclerosis. These findings indicate that location and cell type are key determinants of LOX-1 function, but cell-specific sources of LOX-1-mediated lung protection are unknown. Here, we show that a portion of lung-recruited neutrophils are stimulated to express LOX-1 upon entry into the airspaces of humans and mice during pneumonia. Neutrophil LOX-1 expression correlates with higher levels of chemokine and scavenger receptors, reactive oxygen species, and apoptotic markers that likely influence their phenotype and potentially those of neighboring cells. To determine whether neutrophil LOX-1 contributes to LOX-1-dependent tissue protection, we generated neutrophil-specific LOX-1 knockout mice (LOX-1^ΔMRP8^) and assessed their acute response to two models of bacterial pneumonia, *Escherichia coli* and *Streptococcus pneumoniae.* Elevated metrics of inflammation were detected in LOX-1^ΔMRP8^ mice following both challenges, including reduced survival during *E. coli* infection, without changes in bacterial clearance. Finally, deletion of LOX-1 on neutrophils resulted in modifications to the neutrophil phenotype that may contribute to increased recruitment and inflammatory potential in the airspace. Taken together, LOX-1 is a scavenger receptor that is induced on a population of neutrophils in the pneumonic airspaces, taming further neutrophilic inflammation to limit immunopathology.

## Introduction

Lower respiratory infections are the primary cause of pneumonia, which leads the world in infectious disease-related morbidity and mortality (18 million non-Covid deaths in 2021) [1]. A diverse range of microorganisms, including bacteria, viruses, and fungi, cause pneumonia, challenging the development of broadly applicable therapeutics. Host immunity, while critical for anti-microbial defense, fuels pneumonia-related complications when excessive, increasing the likelihood of acute respiratory distress syndrome, sepsis, disseminated intravascular coagulation, and other causes of pathology. As such, host-directed therapies represent an important potential avenue for clinical intervention. Neutrophils are an essential source of immune protection but are also particularly threatening to tissue integrity during pneumonia, with neutrophil degranulation and activation pathways enriched in patients with severe bacterial pneumonia [2]. Thus, host processes regulating neutrophil function to counter inflammatory injury constitute an important knowledge gap.

While neutrophils are known to promote inflammatory injury due to their specialized roles in pathogen clearance (degranulation, NETosis, reactive species production), their heterogeneity has only recently been appreciated. Through improved flow cytometry techniques and the advent of single cell sequencing, multiple neutrophil subsets have been identified with diverse roles during homeostasis [3], infection [3, 4], autoimmunity [5], and cancer [6, 7]. Specifically in the lung, tumor associated neutrophils vary in their expression of antigen presentation proteins (MHCII, CD74) [8], chemokine receptors (CXCR1, CXCR2, and CXCR4) [9, 10], adhesion markers (CD54) [11] and immunosuppressive proteins (Arg1, PD-L1, IL1RN) [12]. During lung infection, several neutrophil populations have emerged based on their capacity to limit inflammation or promote clearance of pathogens [13], two processes that are not mutually exclusive. For instance, PD-L1+ neutrophils also promote clearance of MRSA and repair liver injury [14]. SiglecF, traditionally thought to be limited to alveolar macrophages and eosinophils, marks a specific subset of neutrophils that promote macrophage expansion and resolution of epithelial damage during acute lung injury [15], but drive the progression of lung emphysema [16] and pulmonary fibrosis [17]. MHCII [18] and PD-L1 [19, 20] have also been identified as markers of lung neutrophils, which are generally thought to be more immunosuppressive, though there are a few exceptions [21].

We have found that hematopoietic-derived LOX-1 prevents inflammatory lung injury during pneumonia, with prominent expression on tissue resident alveolar macrophages and many recruited neutrophils [22]. LOX-1+ neutrophils are transcriptionally unique from their LOX-1-counterparts, enriched in gene expression programs with potential to counteract inflammatory injury [22]. Interestingly, LOX-1 has also been identified as a marker for neutrophil-derived myeloid suppressor cells in cancer [23] and may limit inflammation during SARS-CoV2 infection [24], further supporting LOX-1+ neutrophils as a source of immunomodulation. However, it is insofar unknown whether and how LOX-1 directly plays a role in limiting lung injury and in regulating the neutrophil phenotype. In the current study, we expanded upon our previous work by further defining the expression pattern of LOX-1 on pneumonia-elicited neutrophils in the lungs of humans and mice. Moreover, we determined the direct contributions of neutrophil LOX-1 to their phenotype and overall outcome during pneumonia using a novel conditional knockout mouse model, providing the first evidence that neutrophil LOX-1 directly influences pneumonia outcome.

## Methods

### Sex as a biological variable

Our study included both male and female mice with no differences in phenotype based on sex.

### Rapid Human Autopsy Biobank Samples

Human lung tissues were acquired from the Brain and Body Donation Program of Banner Sun Health Research Institute where retirement-age subjects provided consent for rapid autopsies between the years of 2006-2020, collected from retirement communities with the greater Phoenix, AZ area. The median age of autopsy subjects was 82 years at autopsy and tissues are collected and processed no longer than 3 hours post-mortem, as previously described [25]. For the current work, we compare tissue sections from subjects that died with a pathological diagnosis of pneumonia (and whose H&E-stained lung sections revealed presence of bacteria and abundant neutrophils) to those from control subjects who died without a diagnosis of pneumonia or other lung disease (randomly selected).

### Histology, Immunohistochemistry. & Whole Slide Scanning

Human lung formalin-fixed paraffin-embedded (FFPE) tissues were sectioned at 5 µm using a RM2255 Microtome (Leica, Wetzlar, Germany) for multiplex fluorescent immunohistochemistry (mfIHC). Sections were processed on a Ventana Discovery Ultra autostainer (Roche Diagnostics, Indianapolis, IN, USA) with Benchmark Ultra CC1 antigen retrieval (Tris-based buffer, 95°C, 32 min). After protein blocking with Opal Diluent/Block (Quaternix, Billerica, MA, USA), primary and secondary antibody pairs (ImmPRESS HRP Goat Anti-Rabbit/Anti-Mouse IgG Polymer Detection; Vector Laboratories, Newark, CA, USA; 20 min incubation at 37°C) were developed sequentially with Opal fluorophores (**Table 1**). Nuclear counterstaining was performed with Spectral DAPI (Quaternix) and slides were mounted in Prolong Gold Antifade Mountant (Invitrogen, Waltham, MA, USA). Whole-slide images were acquired on a PhenoImager HTTM Automated Quantitative Pathology Imaging System (Akoya) and spectrally unmixed using InForm v3.0 (Akoya).

**Table 1:**
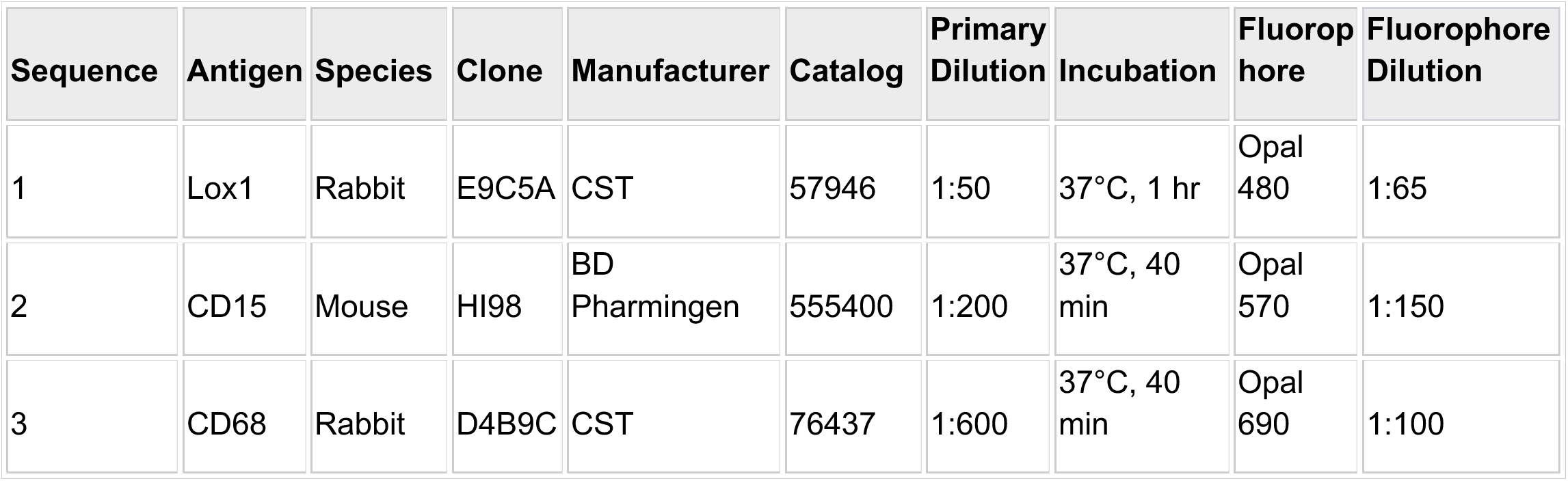

### Whole Slide Image Analysis

Whole slide images of stained lungs were annotated to define a ROI in HALO (v4.2.6399.280, Indica Labs). For quantifying CD15, CD68, or Lox1 immunoreactivity, the HALO Area Quantification (AQ) module (v3.0.3) was fine-tuned based on color and stain intensity to match positive immunoreactivity in the fluorescent images. Phenotypes were defined as CD15+Lox1+, CD15+Lox1−, CD68+Lox1+, and CD68+Lox1−. The algorithm quantified the area of each marker and phenotype within the annotated ROI, outputted in micrometers squared (μm²), and percent area was calculated relative to total area analyzed or total Lox1+ immunoreactivity. Quantitative output was exported as .CSV and analyzed with a two-way ANOVA (condition × phenotype) in Prism.

### Mice

Wild-type C57BL/6J (#000664) and MRP8-Cre-ires/GFP transgenic (#021614) mice were purchased from Jackson Laboratories (Bar Harbor, ME). Olr1-flox (LOX-1-flox) mice were generously provided by Dr. Tatsuya Sawamura (Shinshu University, Japan), where the original mouse line was created and detailed in Supplemental Figure 1. To generate the mice, a target vector was created by first amplifying a 4398 bp-short arm containing exon 5 tagged with SacII and Xhol sites (SacII-13327-17735-Xhol) and inserting an FRT-neo-FRT-loxP cassette followed by Xmal and BanHI sites using pBluescript. A 1928 bp fragment of *olr1* containing exon 6, 7, and an upstream part of exon 8 connected via loxP and tagged with Xmal and BamHI sites was then amplified by PCR from the mouse genome (Xmal-17733-exon 6&7-19661-loxP-BamHI). An IRES2-LacZ cassette (Clontech) with BamHI sites on each end (BamHI-IRES2-LacZ-BamHI) was inserted followed by an 11902bp-long arm containing the remaining part of exon 8 (BamHI-19661-31563-Sall) and a tk cassette. To generate neutrophil-conditional LOX-1 knockout (LOX-1^ΔMRP8^) mice, we bred MRP8-Cre-ires/GFP transgenic mice with homozygous Olr1-flox mice. Heterozygous Cre+ mice and homozygous Cre-littermate controls were used in all experiments. Mice were housed in a specific pathogen free facility at the University of Massachusetts Chan Medical School and provided food and water *ad libitum*. Mice were group-housed (≤5 mice/cage) at room temperature (20°C–23°C) on a 12h light-dark cycle. Both male and female mice were used in experiments and were between 6-16 weeks in age. Unless otherwise specified, all animal outcomes were performed in at least two independent experiments.

### Experimental Pneumonia

Mice were anesthetized using a combination of ketamine (50 mg/kg of body weight) and xylazine (5 mg/kg) administered by intraperitoneal injection (i.p.). A 24-gauge angiocatheter (Becton Dickinson) was inserted into exposed trachea and guided into the left lobe for all intratracheal instillations, as described previously [22]. Approximately 1 x 10^6^ colony forming units (CFU) of *Escherichia coli* (*E*. *coli* serotype 06:K2:H1; ATCC 19138) or *Streptococcus pneumoniae* (serotype 3; ATCC 6303) were delivered as a 50 μl bolus diluted in sterile saline. Mice were humanely euthanized at end points described in results by isoflurane overdose.

### Bone marrow isolation

Bone marrow cells were flushed from the femur and tibia of wild-type mice in cold, sterile PBS, and filtered through a 70 μm filter (Thermo Fisher Scientific). Red blood cells were lysed using a standard RBC lysis buffer (Sigma Aldrich). Neutrophils were subsequently isolated by magnetic separation using a negative selection kit per manufacturer’s protocol (Stem Cell Technologies).

### Bone marrow neutrophil stimulation

Bone marrow neutrophils were isolated using the EasySep™ Mouse Neutrophil Enrichment Kit (Stem Cell™ Technologies) per manufacturer’s protocol. Neutrophils were cultured in 24-well plates, supplemented with RPMI, 10% FBS, 1% Penicillin/Streptomycin, and stimulated with BALF from uninfected mice or mice infected with *E. coli* for 24H (50% BAL collected in complete RPMI + 50% complete RPMI) or 100 ng/ml of ultra-pure lipopolysaccharide from *E. coli* O111:B4 (Sigma Aldrich).

### Bronchoalveolar Lavage

Bronchoalveolar lavage fluid (BALF) was collected post-euthanasia, where an 18-gauge angiocatheter (Becton Dickinson) was inserted into exposed trachea and lungs were washed with 10 total washes of PBS (1 ml/each) and the first 1 ml of lavage was saved separately for protein analyses. Total cell counts were obtained from a LUNA-FL dual fluorescence cell counter (Logos Biosystems). Differential cell counts were determined by flow cytometry as described below. Total protein in BALF was determined by bicinchoninic acid (BCA) assay (MilliporeSigma).

### Cytokine Quantification

BALF cytokines were quantified using a mouse magnetic 23-plex, customized multiplex cytokine panel (Promega, ProcartaPlex) per manufacturer’s instructions.

### Bacteriology

Blood CFU was measured in heparanized blood collected post-euthanasia from the inferior vena cava. Blood was diluted in sterile PBS (1:10-1:1000) and plated at 100 μl on sheep blood agar plates (Carolina Biological Supply) and cultured at 37°C, 5% CO2 overnight, and enumerated the following day. Blood CFU was calculated as CFU/ml. Lung CFU was determined by immediately homogenizing infected lobes using a Bullet Blender (Next Advance) in sterile distilled water and performing serial dilutions. After overnight incubation at 37°C, 5% CO2, colony numbers were counted and CFU calculated as total CFU per lung.

### Collagenase-Digested Lung Processing

Single cell suspensions were generated by enzymatic digestion of manually minced lung tissue with collagenase digestion solution (1× PBS, 1 mg/mL type II collagenase from Worthington Biochemicals, 150 μg/mL DNase from MilliporeSigma, 2.5 mM CaCl2) at 37°C, 200 rpm, 1 hour. Digested tissue was then passed through a 70 μM filter. The filter was then washed with an additional 2-3 ml of PBS to remove extra cells. After pelleting cells at 300 x g for 5 minutes, red blood cells were lysed using a standard RBC lysis buffer per manufacturer’s protocol. Cells were enumerated with LUNA-FL dual fluorescence cell counter (Logos Biosystems) and ∼1 x 10^6^ cells were stained for flow cytometry.

### Intravascular and extravascular discrimination of CD45+ cells

To separate intra- and extravascular cells for flow cytometry, we administered 100 μl α-mouse, α-CD45.2 antibody (diluted 1:10 in sterile saline) 3-5 minutes prior to euthanasia by retroorbital injection. To determine intravascular CD45 positivity, we used an additional mouse (treated similarly – e.g. infected for 24H) that did not receive intravenous CD45 antibody as our FMO (fluorescence minus one), unstained, green fluorescence protein (GFP), and live/dead single stain control.

### Flow Cytometry

After generating the single cell suspension at 1 x 10^6^ cells, we stained for viability in 100 ul 1X PBS plus Zombie-UV or Zombie-NIR (1:300 dilution) for 20 minutes at room temperature, in the dark. After washing cells 1X, they were resuspended in Fc receptor blockade (FcBlock, eBioscience) and stained with the appropriate cocktail of monoclonal antibodies (**Table 2**) for 30 minutes, on ice, in the dark. After a 1X wash step, cells were resuspended in FACS buffer (1X PBS, 0.6% fetal bovine serum, 0.4% EDTA). Flow cytometry data were acquired using a 5-laser, 64-channel Cytek Aurora.

**Table 2.**
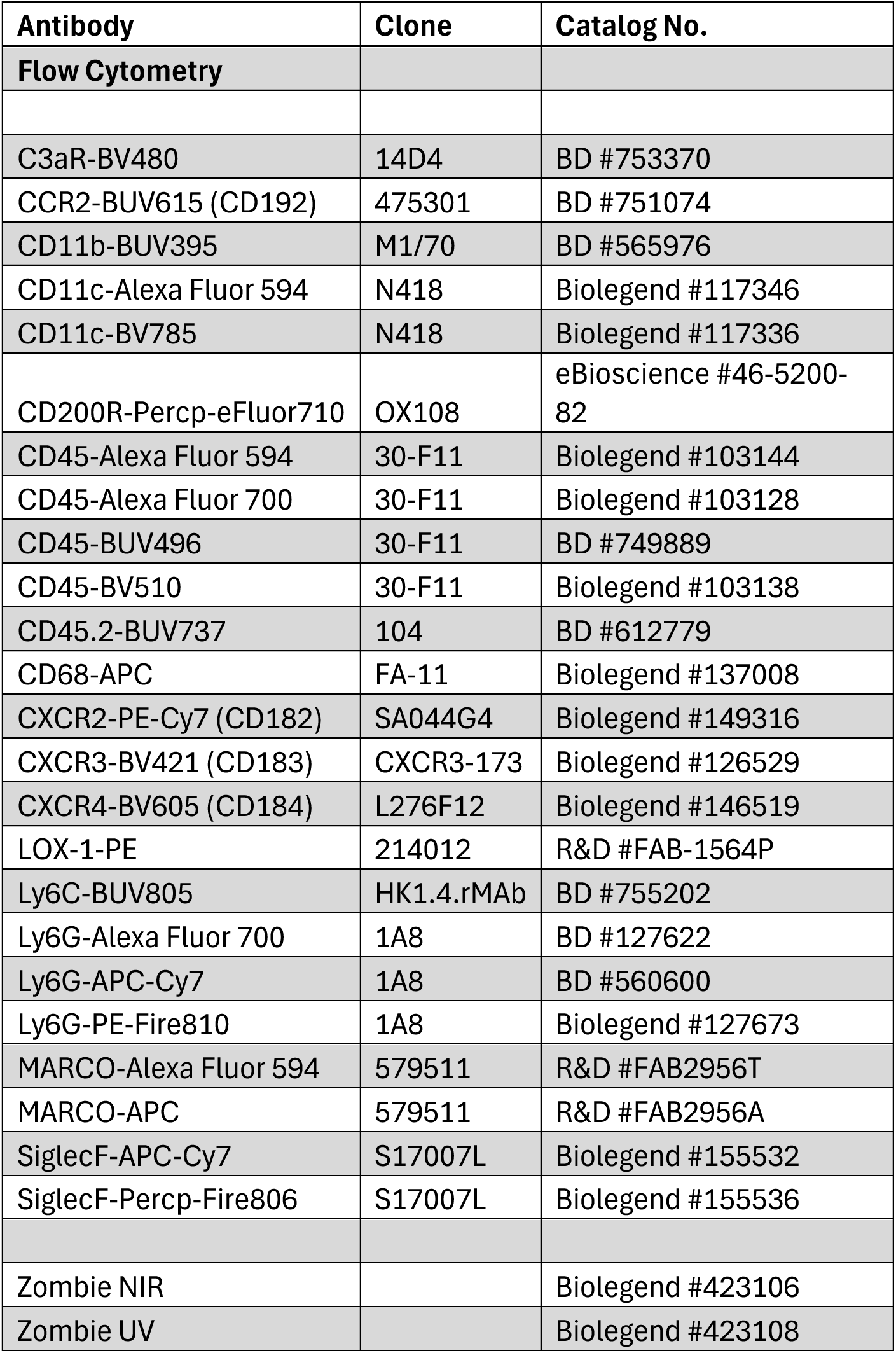

### Statistics

Statistical analyses were performed in Prism V.10 (GraphPad Software). Data are presented as mean ± SEM. Sample size in each experiment is described within the figure legend where n = number of mice or human subjects. Data points are indicative of biological replicates, except for survival curves, which are displayed as summary data. Normality of data was determined using a Shapiro-Wilk test. Data that were not normally distributed were either analyzed using nonparametric tests or were log-transformed prior to analysis. Statistical tests used are described in the figure legends, with two group comparisons made with two-tailed t-tests and comparisons with more than two groups with analysis of variance (ANOVA). Multiple comparison’s testing was performed post-hoc ANOVA analyses. Differences were considered significant if P < 0.05, and the following represents the level of significance: ****P < 0.0001, ***P < 0.001, **P < 0.01, and *P < 0.05.

### Study Approval

Animal procedures were approved by the Institutional Care and Use Committee at UMass Chan Medical School (protocol no. PROTO202100149) and by the Institutional Care and Use Committee at the University of Iowa Carver College of Medicine (protocol no. 4062599). Studies involving human participants were approved by the Banner Sun Health Research Institute. Written informed consent by each patient or legal proxy for medical decision was a requirement for study inclusion.

## Results

### Human neutrophils heterogeneously express LOX-1 in lung tissue that is independent of infection status

We used lung samples from human subjects who died with or without pneumonia after consenting for rapid autopsy [25] to determine whether and which cells were LOX-1 sources in infected human lungs. Lung sections were assessed by immunofluorescent staining and probed for LOX-1 (cyan) along with CD68 (red) and CD15 (yellow) as markers of macrophages and neutrophils, respectively (**Fig 1**). Macrophage LOX-1 expression was lower than expected, albeit detectable (highlighted with white arrows), in both control and pneumonic samples (∼1-4.5%), given previous data generated in our lab [22] and online data resources revealing extremely prominent LOX-1 mRNA in alveolar macrophages. However, macrophage LOX-1 was significantly increased during pneumonia, consistent with our prior mouse studies (**Fig 1A**) [22]. Neutrophils exhibited a high, but heterogeneous expression of LOX-1, consistent with our prior observations [22]. Depending on the patient sample, we measured between 26-78% LOX-1+ neutrophils (**Fig 1B**), with no significant differences detected between control and pneumonia samples, though LOX-1 expression was numerically elevated when correcting for total cells (**Fig 1B**). Ultimately, these findings confirm LOX-1 expression on human lung neutrophils.

**Figure 1:**
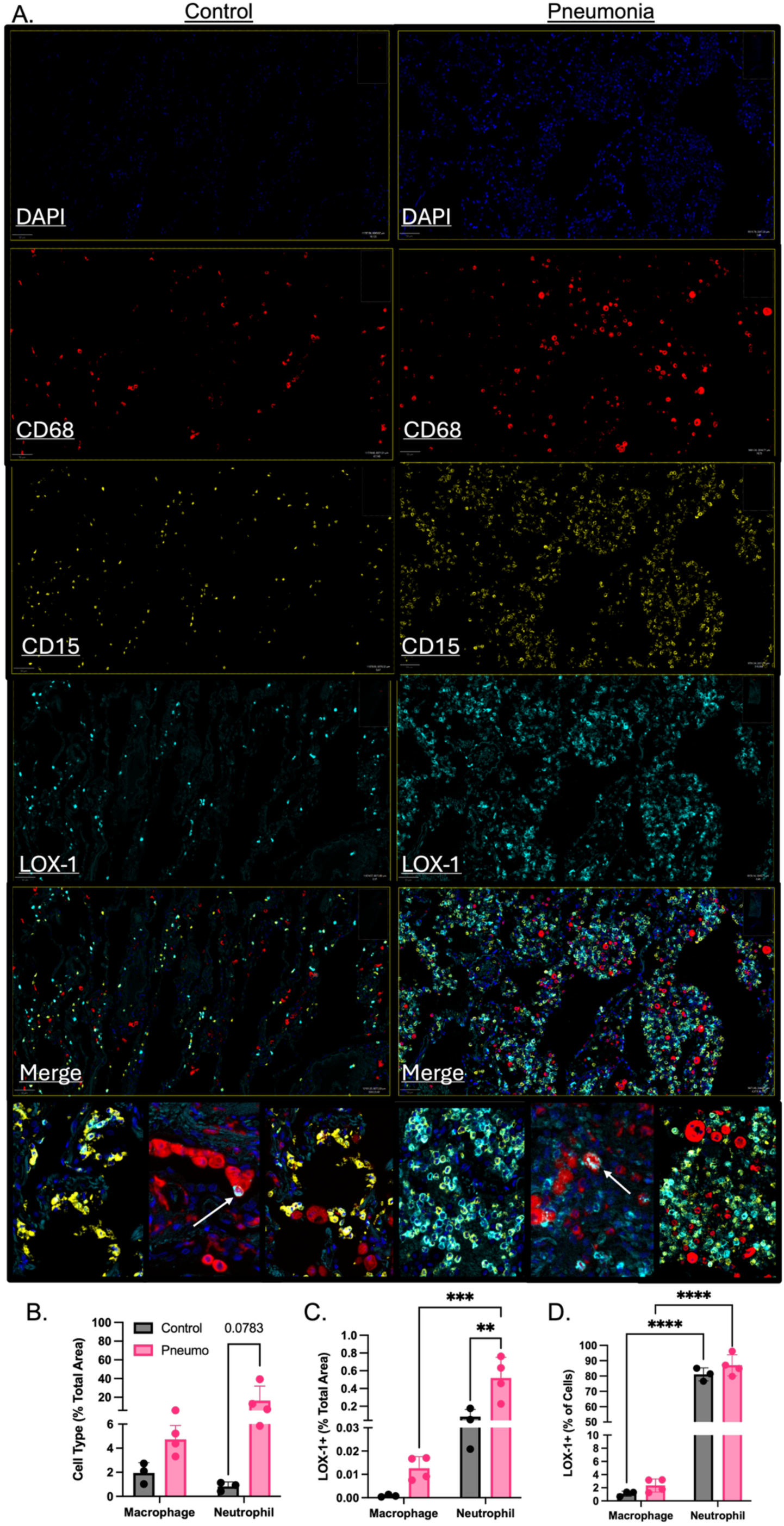
Expression of LOX-1 is detectable on human lung neutrophils regardless of infection status. A) Rapid autopsies were performed on control or pneumonic human subjects. Immunofluorescence was used to detect LOX-1 (cyan), CD15 (neutrophils; yellow) and CD68 (macrophages; red) on lung sections with DAPI staining nuclei in blue. B) Percent of macrophages and neutrophils were determined against total area (%CD68+ or %CD15+/area analyzed). C) LOX-1 expression was determined on CD68+ cells as % LOX-1+/CD68+ (macrophages) and on CD15+ cells as % LOX-1+/CD15+ (neutrophils) D) LOX-1 positivity by cell type was determined on CD68+ and CD15+ cells as % LOX-1+ out of all CD68+ (macrophage) cells % LOX-1+ out of all CD15+ (neutrophil) cells. Arrows highlight CD68/LOX-1 co-expression. *p<0.05 and ***p<0.001 for Welch’s t-test.

### Stimulation of bone-marrow neutrophils with pneumonic bronchoalveolar lavage fluid drives expression of LOX-1

Given the consistency of neutrophil LOX-1 expression in humans to our previous results, where we showed substantial heterogeneity in airspace neutrophil LOX-1 expression using *E. coli* as an infection model [22], we then determined whether LOX-1 heterogeneity is inherent to neutrophils at baseline, or rather, a consequence of exposure to the airway environment. To test this, we isolated naïve bone marrow-derived neutrophils or circulating neutrophils from heparinized blood following infection with *E. coli* (circulating cells). Neutrophils outside of the lung environment exhibited markedly lower levels of LOX-1 expression (**Fig 2A-B; BALF vs Blood**). Moreover, stimulation of bone marrow-derived neutrophils with cell-free and antibiotic-treated bronchoalveolar lavage fluid (BALF) from *E. coli*-infected, but not uninfected mice, significantly induced LOX-1 expression on WT neutrophils at a level that is akin to mice *in vivo* and human subjects (**Fig 2C-D**), while neutrophils treated with media alone or uninfected BAL did not exhibit LOX-1 expression that was substantially higher than KO cells (**Fig 2C-D**). To confirm whether neutrophil heterogeneity is relevant to other infection models, we also measured LOX-1 surface expression on neutrophils isolated from the BALF of mice infected with *Streptococcus pneumoniae* (serotype 3) from 12H to 48H post-infection. Like *E. coli* infection, during *S. pneumoniae* infection, we detect a substantial increase in LOX-1+ neutrophils with nearly 100% of BAL neutrophils expressing LOX-1 at 12h post-infection, dropping to 64% and 61%, respectively, at 24 and 48h-post-infection (**Supplemental Fig 2**), suggesting that LOX-1 expression on later waves of neutrophils may be reduced due to differences in the life cycle (emergency granulopoiesis) or the environment, wherein secreted and intramembrane proteases may elicit LOX-1 cleavage from the cell surface [26, 27].

**Figure 2:**
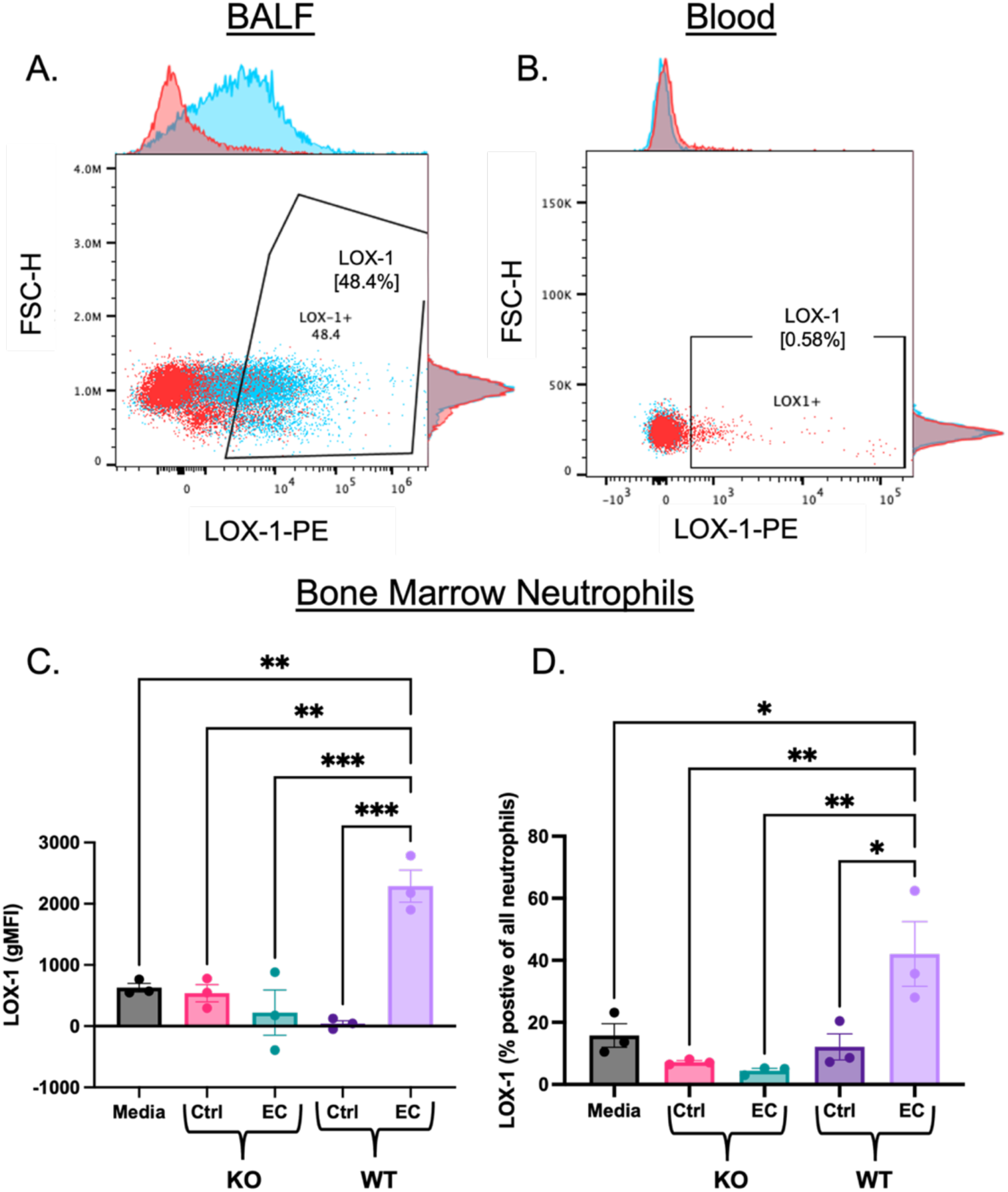
Neutrophils increase expression of LOX-1 when exposed to the infected airspace. A) Male and female C57BL/6 mice were infected with 1x10^6^ CFU *Escherichia coli* for 24H. LOX-1 surface expression was measured by flow cytometry on neutrophils (Live/CD45+/CD11b+/Ly6G+) isolated from the A) bronchoalveolar lavage fluid (BALF) and B) heparinized blood. D) Bone marrow neutrophils were isolated from uninfected C57BL/6 and full body LOX-1 knockout (KO) mice and stimulated with media, uninfected BALF or BALF collected from mice infected with *E. coli* for 24H. LOX-1 surface expression was similarly measured at 24H post stimulation *ex vivo*. *p<0.05, **p<0.01, and ***p<0.001 one-way ANOVA with Tukey’s multiple comparisons. Representative histograms show LOX-1 expression in sample (blue) and fluorescence minus one (FMO) control (red).

### LOX-1+ neutrophils are phenotypically distinct

Given the heterogeneity found in neutrophil LOX-1 expression, we next evaluated immunophenotypic differences between LOX-1- and LOX-1+ neutrophils, including receptors linked to neutrophil chemotaxis and activation. Wild type mice were infected with *E. coli* for 24h and LOX-1+ and LOX-1-cells were discriminated by flow cytometry (LOX-1 positivity determined vs. fluorescence minus one control) (**Fig 3A**). Interestingly, several receptors associated with neutrophil retention, transmigration, and chemotaxis were significantly increased in LOX-1+ neutrophils, including CCR2, CXCR3, and CXCR4 (**Fig 3B**). Similar results were measured in mice infected with *S. pneumoniae*, albeit with minor differences (**Supplemental Fig 2**). After *S. pneumoniae* infection, LOX-1+ neutrophils also had increased CXCR2 expression (**Supplemental Fig 3**), which is the primary chemokine receptor known to promote neutrophil recruitment. These findings indicate that LOX-1 expression is tethered to a distinct neutrophil phenotype.

**Figure 3:**
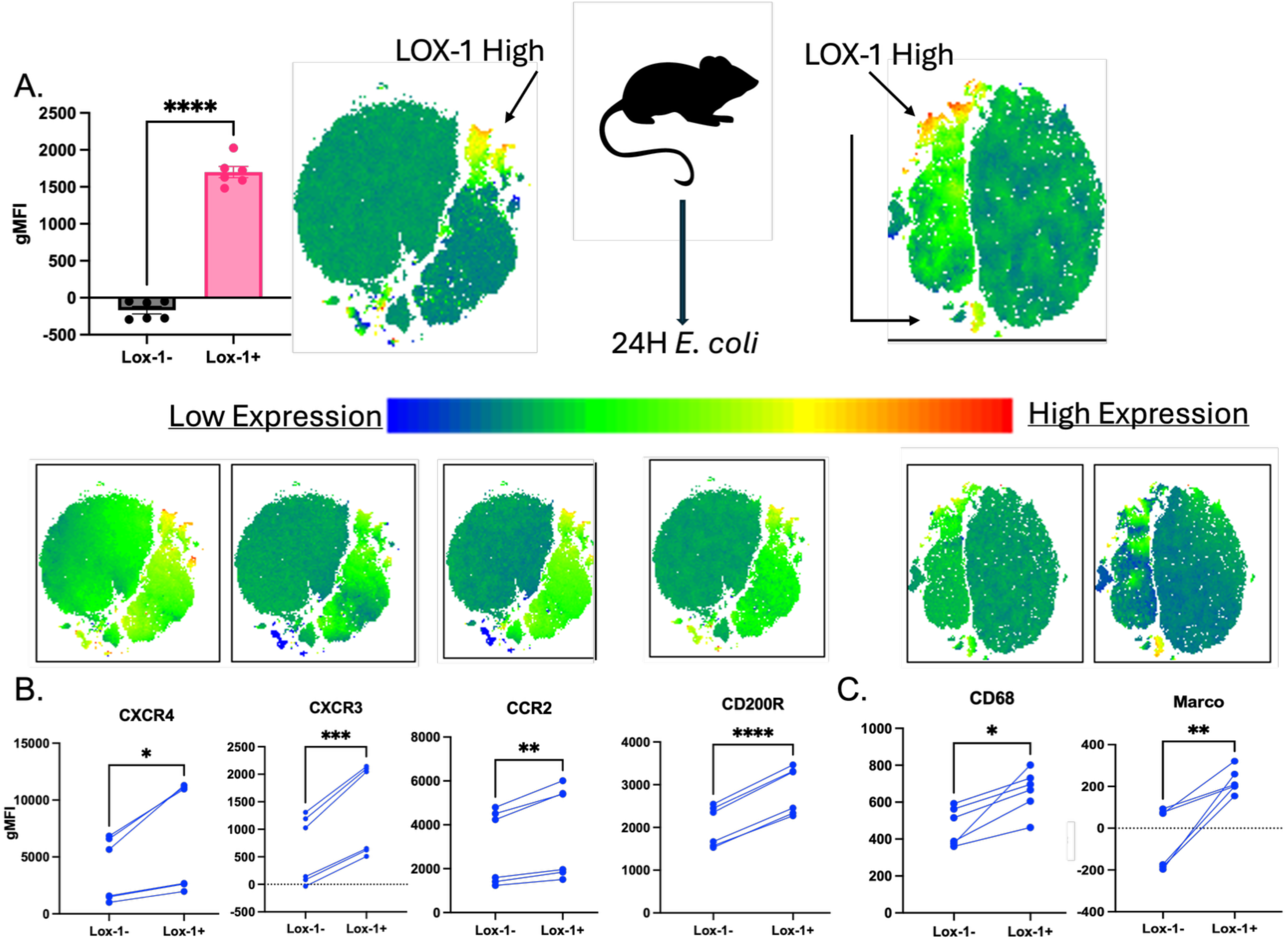
LOX-1+ neutrophils are phenotypically distinct from LOX-1-cells. A) Male and female C57BL/6 mice were infected with 1x10^6^ CFU *Escherichia coli* for 24H. LOX-1 surface expression was measured by flow cytometry on neutrophils (Live/CD45+/CD11b+/Ly6G+) isolated from bronchoalveolar lavage fluid (BALF) and fluorescence minus one (FMO) control was used to determine LOX-1 positivity. LOX-1 expression was quantified with PE gMFI and visually shown in yellow-red (high LOX-1) in tSNEs from two separate flow cytometry experiments. B) Chemokine (CXCR4, CXCR3, CCR2) and C) scavenger receptors (CD200R, CD68, Marco) were then assessed by flow cytometry on LOX-1+ and LOX-1-neutrophils using gMFI for quantification and visually represented in tSNEs above each receptor panel. *p<0.05, **p<0.01, ***p<0.001, ****p<0.0001 paired t-test.

### LOX-1+ neutrophils express higher levels of macrophage-associated scavenger receptors

Scavenger receptors, such as Marco and CD36, have been studied extensively in macrophages with reported roles in promoting phagocytosis [28], lipid uptake [29], and clearance of apoptotic/necrotic cells and debris [30]. However, the role of scavenger receptors on neutrophils is less well-characterized and has focused mainly on tumor-associated neutrophils. Previous RNA sequencing data from our group showed that LOX-1+ neutrophils exhibit increased expression of *Marco* and *CD68* (15-fold and 1.45-fold, respectively, FDR<0.05) ([22]). To determine whether scavenger receptors are differentially expressed at the protein level, we infected WT mice for 24h with *E. coli* and separated BALF neutrophils as described above and measured the expression of Marco, CD68, and CD200R, among a larger panel of neutrophil lineage (Ly6G, CD11b) and functional markers (described above), by flow cytometry. Consistent with transcriptional data, we found that LOX-1+ neutrophils exhibit measurable levels of both receptors and significantly higher levels of Marco and CD68 in LOX-1+ cells (**Fig 3C**, 6-fold and 1.4-fold, p<0.05). In addition, CD200R is an immunosuppressive receptor that interacts with CD200. In neutrophils it has been shown to drive both immunosuppression and antimicrobial activity. Here, LOX-1+ neutrophils have approximately 1.4-fold higher levels of CD200R expression during *E. coli* infection (**Fig 3C**), consistent with their immunoregulatory potential [22]. Like chemokine receptors, we detected a significant and similar increase in Marco following *S. pneumoniae* infection (**Supplemental Fig 3**). Finally, SiglecF has been identified on subsets of neutrophils that exhibit unique, tumor-promoting properties in the lung [31]. Consistent with data revealing LOX-1 as a biomarker for tumor-infiltrating PMN-MDSCs [23] and our own findings suggesting a less inflammatory profile in LOX-1+ neutrophils [22] we measured an elevated level of SiglecF surface expression in LOX-1+ neutrophils during *S. pneumoniae* infection (**Supplemental Fig 3**).

### LOX-1+ neutrophils are enriched for apoptosis and ROS production

Efferocytosis is a mechanism for the clearance of cellular debris that leads to decreased inflammation through removal of potentially necrotic cells and direct changes on macrophage activation. These anti-inflammatory processes are triggered by apoptosis. Interestingly, transcriptional profiling of LOX-1+ versus LOX-1-neutrophils showed that *Anxa5* (annexin A5) is upregulated on LOX-1+ neutrophils 24H post-infection with *E. coli* [22], prompting us to compare apoptosis between these two cell populations at 18H and 24H post-*E. coli* infection. LOX-1+ cells exhibit higher levels of early apoptotic cell markers (Annexin-5+/PI-, **Fig 4A**) at both time points post infection, with higher overall levels of apoptosis at 24H post-infection as expected. Thus, LOX-1+ neutrophils may be predisposed to apoptosis, increasing the likelihood of resolving effects from efferocytosis. Since LOX-1 is known to activate NADPH oxidase, a primary mechanism for generation of reactive oxygen species (ROS) in neutrophils, we similarly measured total ROS production at 24H post infection with *E. coli* in LOX-1+ and LOX-1-neutrophils by flow cytometry. In line with previous reports, LOX-1 expression correlated with significantly higher levels of total ROS production (**Fig 4B**), indicating LOX-1 may promote both anti-microbial and anti-inflammatory functions.

**Figure 4:**
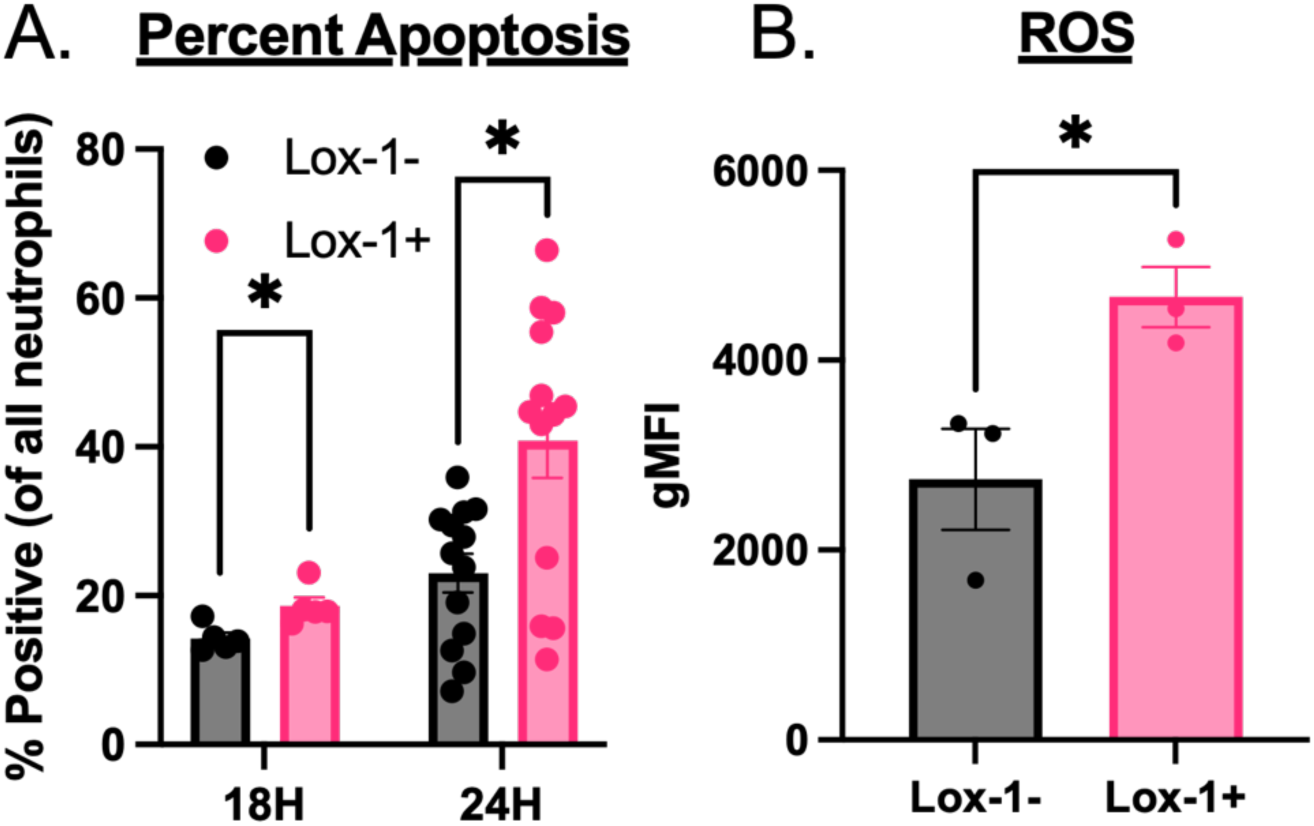
LOX-1+ neutrophils are more apoptotic and have elevated ROS. A) Male and female C57BL/6 mice were infected with 1x10^6^ CFU *Escherichia coli* for 18-24H. LOX-1 surface expression was measured by flow cytometry on neutrophils (Live/CD45+/CD11b+/Ly6G+) isolated from bronchoalveolar lavage fluid (BALF) and fluorescence minus one (FMO) control was used to determine LOX-1 positivity. Early apoptosis was measured with Annexin-5 and propidium iodine (PI) staining where early apoptotic cells are Annexin-5+ and PI. Percent apoptosis was assessed as % of all neutrophils. B) Reactive oxygen species were measured by flow cytometry at 24H post infection with *E. coli.* *p<0.05 for paired t-test.

### Neutrophils are the primary source of LOX-1 in pneumonic airspaces

Given the suggested role for neutrophil LOX-1 in promoting LOX-1-dependent lung protection during pneumonia [22], we generated a neutrophil conditional LOX-1 knockout mouse by crossing mice with homozygous floxed *Olr1*. Homozygous floxed *Olr1* mice, produced in collaboration with Dr. Tatsuya Sawamura (Shinshu University), were generated via targeting vectors directed to insert loxP sites around exon 6 of the *olr1* locus (described in Methods; **Fig 5A**). Next, we crossed mice containing homozygous floxed alleles with those expressing Cre-recombinase under control of the Mrp8 promoter to specifically target neutrophils. To determine whether Cre-mediated deletion of LOX-1 on neutrophils was successful using this genetic approach, we infected LOX-1^ΔMRP8^ mice (Cre+) and Cre-littermate controls with *E. coli* for 12-48h. We previously showed that bone marrow chimeras generated from WT and full body LOX-1 knockout mice have significantly lower levels of soluble LOX-1 (sLOX-1) in BALF that is likely neutrophil-derived [22] through proteolytic cleavage. Here we similarly measured sLOX-1 in BALF of Cre+ and Cre-mice at both 12 and 24H post-infection and measured significantly reduced sLOX-1 concentrations in Cre+ mice (**Fig 5B**). Moreover, surface expression of LOX-1 is significantly reduced in Cre+ mice to comparable levels of full body LOX-1 knockout mice (**Fig 5C**), while LOX-1 expression remained unchanged in alveolar macrophages (data not shown).

**Figure 5:**
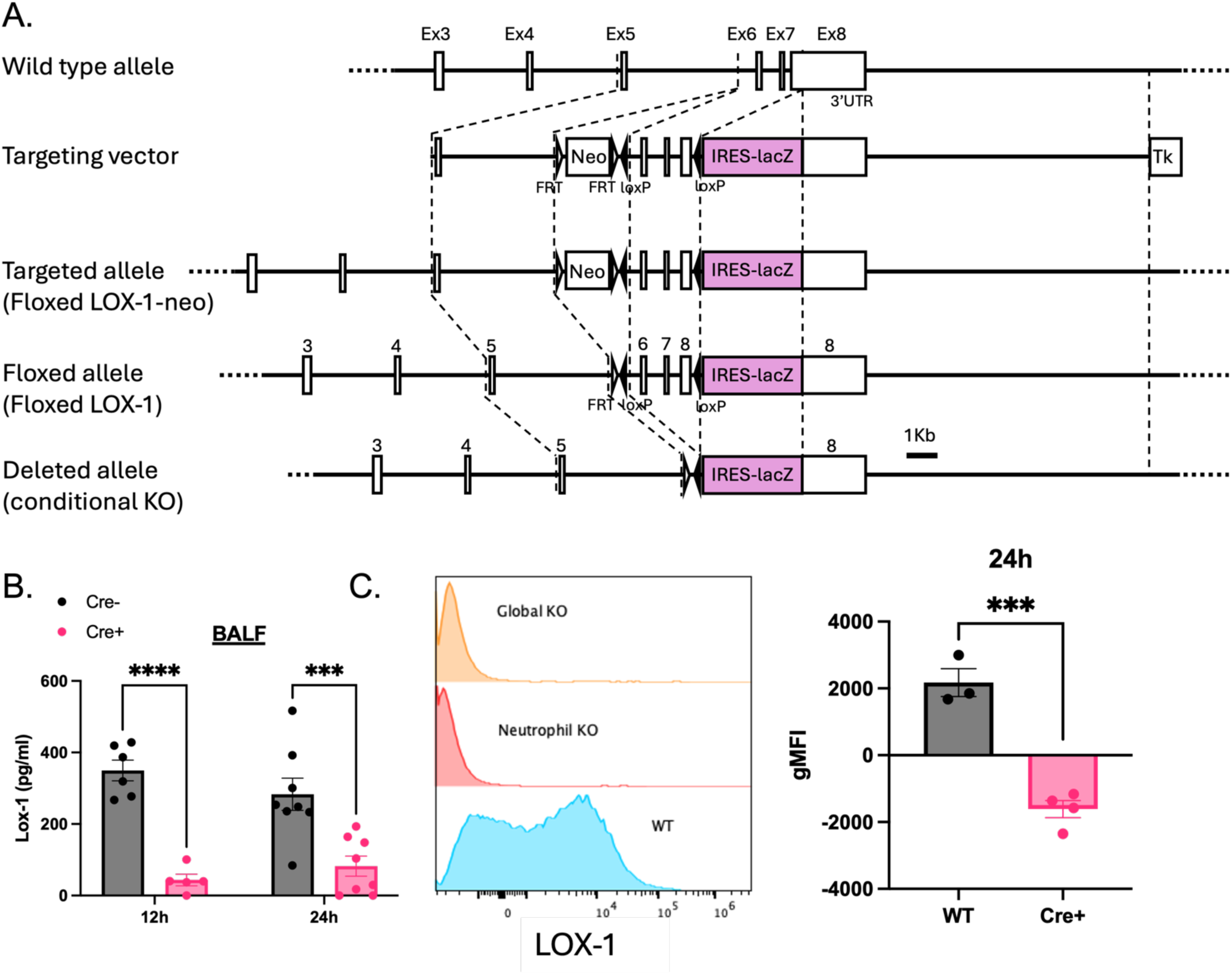
Generation of conditional neutrophil knockout mouse containing loxP sites surrounding critical exons for LOX-1. A) A targeting vector was generated to introduce loxP sites around exons 6,7, and an upstream part of exon 8 using the pBluescript phagemid and a LacZ cassette to identify successful clones that were subsequently used to transfect mouse embryonic stem cells. Mice were subsequently crossed with heterozygous Cre-expressing mice under an Mrp8 promoter. B) Cre+ and Cre-LOX-1^ΔMRP8^ mice were infected with 1x10^6^ CFU of *Escherichia coli* for 12-24H and soluble LOX-1 levels (pg/ml) were determined in BALF at 12 and 24H post-infection. C) LOX-1 surface expression was measured on all neutrophils (Live/CD45+/Ly6G+/CD11b+) at 24H post-infection.

### Neutrophil LOX-1 expression reduces inflammatory injury and improves survival during *E. coli*-induced pneumonia

We previously showed that intratracheal neutralization of LOX-1 promotes inflammation, injury, and mortality during *E. coli* infection and that LOX-1-dependent protection was at least partially mediated by hematopoietic cells [22]. To test whether deletion of LOX-1 on neutrophils influences the severity of *E. coli-*induced pneumonia, we infected both Cre+ and Cre-mice for 12-48H. Mortality was significantly increased by 30H post-infection (**Fig 6A**), similar to mice that received intratracheal neutralization of LOX-1 prior to infection [22]. Given that *E. coli* is typically self-limiting in the lungs of mice, along with our previously described role for LOX-1 in tissue protection [22], we then compared total BALF protein levels as a metric of proteinaceous edema. BALF total protein concentrations were significantly elevated in Cre+ mice at 30H post-infection, corresponding with reduced survival (**Fig 6B**). Finally, we performed flow cytometry on both BAL and lung tissue cells to determine differences in extra-vascular cell number and phenotype. We did not detect differences in neutrophil numbers (data not shown), but alveolar macrophages were significantly elevated in Cre+ mice at 12H post-infection (**Fig 6C**) with *E. coli*. Increased AMs correlated with an increase in extravascular BAL and tissue monocytes (**Fig 6D-E**), possible serving as a source of the former through macrophage replacement. Inflammatory monocytes could be a cause or consequence of increased inflammation. In line with this, tissue and BAL monocytes from Cre+ mice had significantly higher expression of CD11b, an integrin that is increased on activated monocytes, promoting their adhesion and recruitment (**Fig 6D-E**). Notably, macrophage numbers were equivalent between groups at baseline, indicating that changes are infection-dependent and not developmental (data not shown).

**Figure 6:**
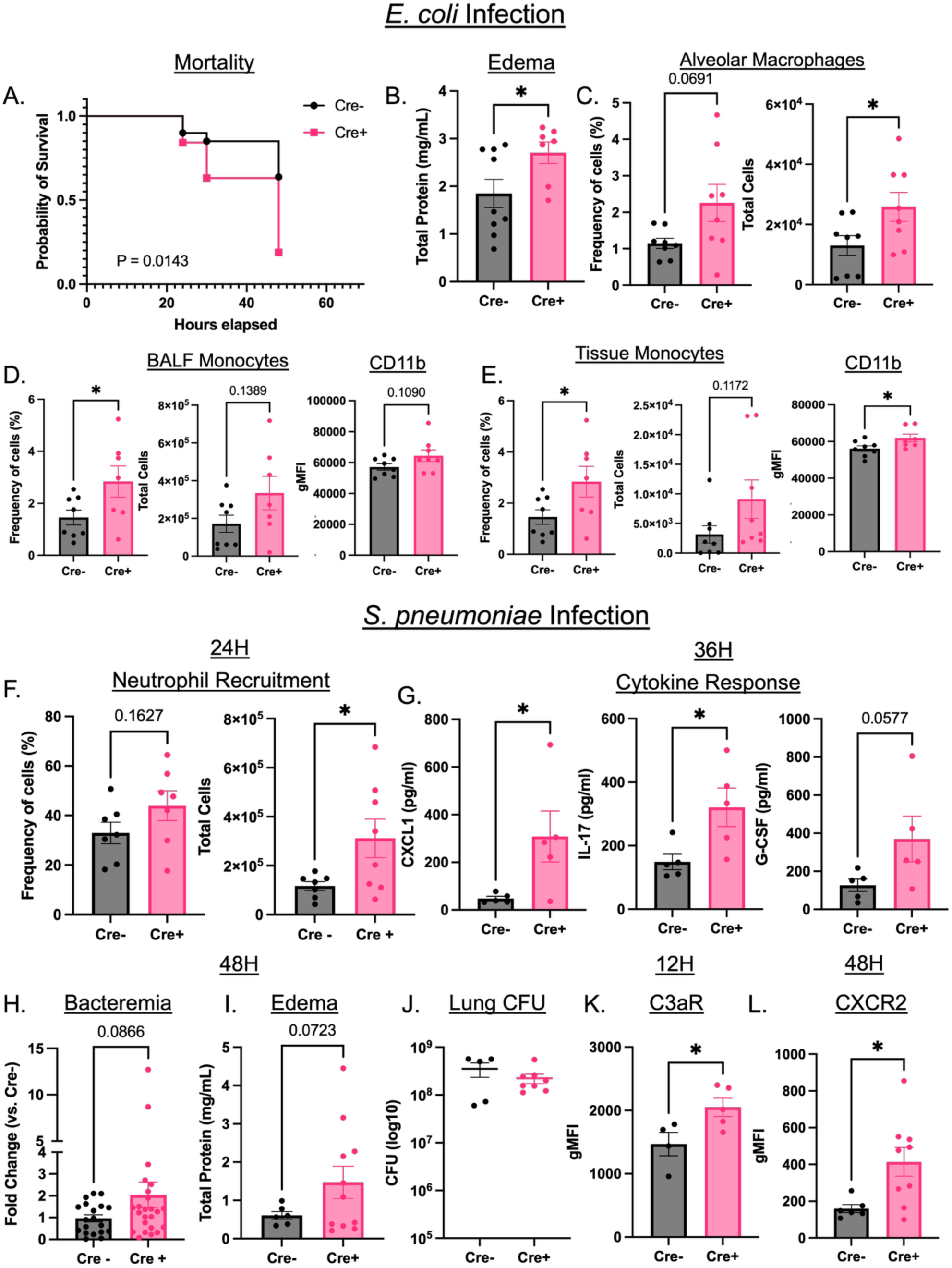
Conditional deletion of LOX-1 in neutrophils exacerbates inflammatory injury and death during pneumonia. A) Cre+ and Cre-LOX-1^ΔMRP8^ mice were infected with 1x10^6^ CFU of *Escherichia coli* for 12-48H and survival was assessed up to 48H post-infection. B) total protein (mg/ml) was measured as a surrogate of alveolar edema at 30H post-infection (to limit survival bias). C-E) BALF alveolar macrophages (Live/CD45+/CD11c+/SiglecF+) and extravascular tissue (intravenous anti-CD45.2-) inflammatory monocytes (Live/CD45+/CD11b+/Ly6C+/Ly6G-) frequency and numbers were measured at 12H post-infection along with CD11b expression on D) BALF and E) tissue monocytes. F) Cre+ and Cre-LOX-1^ΔMRP8^ mice were infected with 1x10^6^ CFU of *S. pneumoniae* (serotype 3) for 12-48H and total BAL neutrophil (Live/CD45+/Ly6G+/CD11b+) numbers and frequency were determined at 24H post-infection along with G) BAL cytokines CXCL1, IL-17, and G-CSF at 36H post-infection. H) Blood bacteria, I) BALF total protein, and J) total lung CFU were determined at 48H post-infection. Frequency of all cells was determined as % of all live cells and backcalculated to the total number of cells. K) C3aR (12H) and CXCR2 (48H) surface expression was measured on extravascular (i.v. CD45.2-) lung neutrophils (Live/CD45+/Ly6G+/CD11b+) collected from collagenase-digested lung tissue. *p<0.05, ***p<0.001, and ****p<0.0001 for student’s t-tailed or Welch’s t-test. Survival was measured with Log-rank (Mantel Cox) test.

### Neutrophil LOX-1 limits inflammatory consequences of pneumococcal pneumonia

Previously, we showed intratracheal neutralization of LOX-1 also leads to increased inflammatory injury during *Streptococcus pneumoniae*-induced pneumonia (serotype 3), though protection is more subtle, as *S. pneumoniae* does not elicit the same degree of alveolar edema at 24H [22]. Given that neutrophil LOX-1 is protective against *E. coli*-induced pneumonia (**Fig 6A-E**), we next determined whether this finding was also applicable to *Streptococcus pneumoniae*. In contrast to *E. coli*, we did not measure any increase in mortality or monocyte or macrophage numbers (data not shown), but total BAL neutrophils were significantly elevated at 24H post infection (**Fig 6F)**; Notably, CXCL1 and IL-17 were significantly elevated at 36H post-infection, with a trending (P=0.0577) increase in G-CSF (**Fig 6G**). IL-17 is a critical cytokine that limits infection of *S. pneumoniae* [32], eliciting neutrophil recruitment through induction of chemokines such as CXCL1, -2, and -5 [33-35], and G-CSF to promote emergency granulopoiesis [36]. Altogether, these cytokines and chemokines may contribute to the significant increase neutrophil recruitment measured in Cre+ mice. Akin to intratracheal LOX-1 neutralization [22], we measured a trend towards increased alveolar edema (P=0.07) and bacteremia (P=0.09) in Cre+ mice (**Fig 6H-I**) with no difference in bacterial clearance (**Fig 6J**), indicating that LOX-1+ neutrophils may be protective against lung injury at later time points post-infection as seen with *E. coli*. Given that neutrophil LOX-1 deletion led to increased neutrophil recruitment (**Figs 6F**), we measured chemokine receptors expressed on neutrophils that may contribute to increased chemotaxis and inflammatory injury. Spectral flow cytometry was employed to measure cell surface markers on extra-vascular neutrophils at 12-24H post-infection with *E. coli* and 12-48H post-infection with *S. pneumoniae*. Notably, at 12H post-infection we measured a significant increase in the expression of neutrophil C3aR, a receptor that responds to the complement protein, C3a, during both models of lung infection (**Supplemental Fig 4; Fig 6K**). Cre+ mice also expressed significantly higher CXCR2 at 48H post-infection with *S. pneumoniae* in both intra (data not shown)- and extravascular cells (**Fig 6L**), possibly contributing to alterations in neutrophil recruitment (**Fig 6F).**

### LOX-1 regulates expression of CXCR4 and other scavenger receptors

Given the substantive receptor differences detected in LOX-1+ vs LOX-1-neutrophils (**Fig 3; Supplemental Fig 2**), we also measured the expression of LOX-1-associated receptors in Cre+ vs Cre-mice during infection with *S. pneumoniae* using spectral flow cytometry. Consistent with our findings above that LOX-1+ neutrophils express higher levels of CXCR4 vs LOX-1-cells (**Fig 3B & Supplemental Fig 1**), Cre+ mice exhibited significantly reduced expression of CXCR4 at 24H post-infection (**Fig 7A**). Moreover, neutrophil LOX-1 deletion led to a significant reduction in CD68 expression (**Fig 7B**) and, surprisingly, significantly higher Marco expression in Cre+ mice at 24H post-infection with *S. pneumoniae* (**Fig 7C**), further indicating that LOX-1 regulates neutrophil phenotype, potentially leading to differences in pneumonia severity.

**Figure 7:**
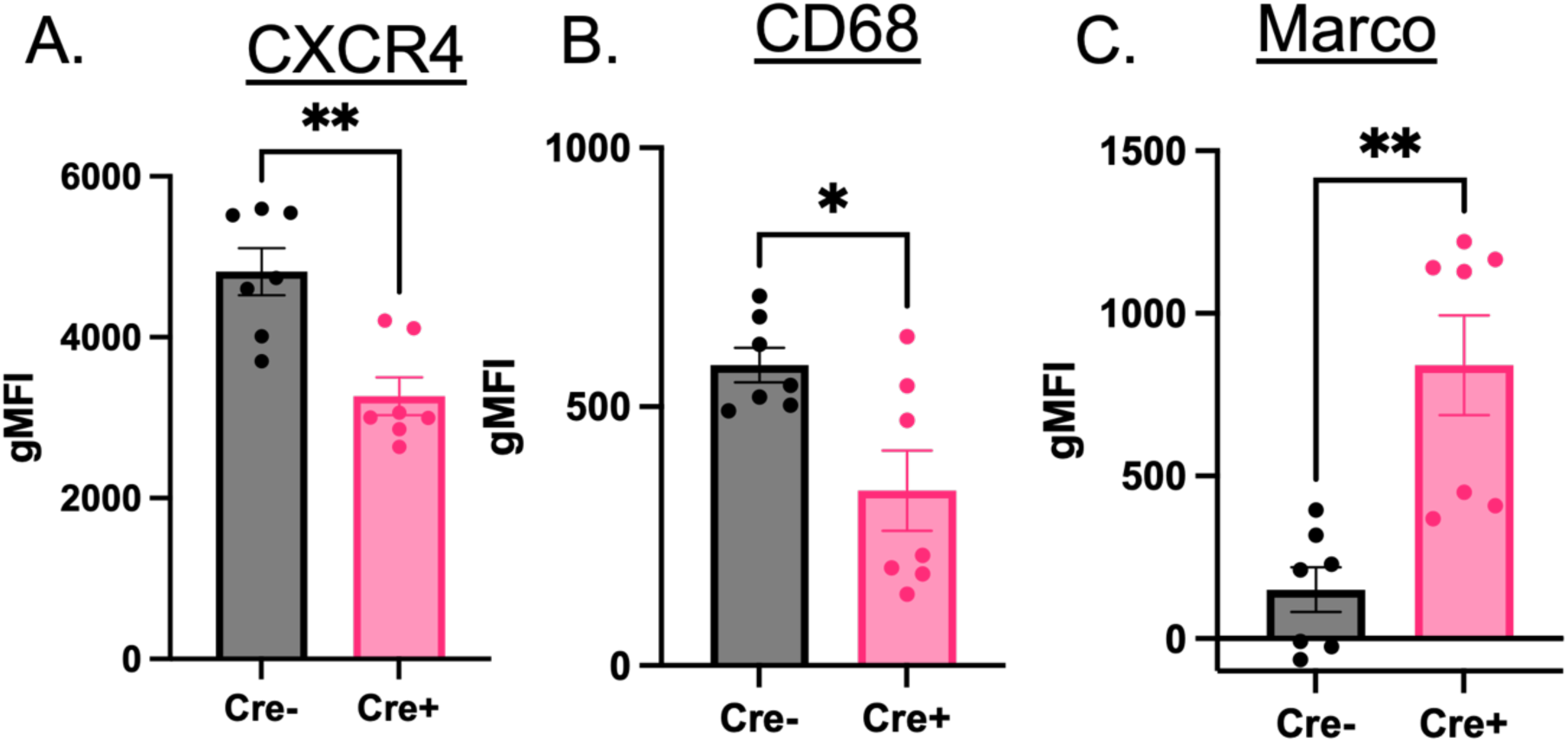
Conditional deletion of LOX-1 alters neutrophil phenotype during lung infection. Cre+ and Cre-LOX-1^ΔMRP8^ mice were infected with 1x10^6^ CFU of *Streptococcus pneumoniae* (serotype 3) for 24H. BALF neutrophils (Live/CD45+/Ly6G+/CD11b+) were isolated and assessed for expression of A) CXCR4, B) CD68, and C) MARCO by flow cytometry. **\***p<0.05 and **p<01 for student’s t-tailed or Welch’s t-test.

## Discussion

In previous work, we identified LOX-1 as a novel, protective receptor against lung injury during pneumonia, with no major impact on bacterial clearance [22]. Protective properties of LOX-1 were determined to result from neutrophil and macrophage LOX-1 expression, with LOX-1+ neutrophils exhibiting transcriptional upregulation of genes that regulate cholesterol metabolism and inflammation [22]. Given the broad expression of LOX-1 on lung cells, we were previously unable to specifically determine the role of neutrophil LOX-1 during respiratory infection. In the current study, we generated a neutrophil-specific LOX-1 knockout mouse (LOX-1^ΔMRP8^) using the Cre-lox system (**Fig 5**), with the help of our collaborators, that do not bear any major differences in growth and development, nor do they exhibit obvious immune dysfunction at baseline. In response to lung infection, however, LOX-1^ΔMRP8^ (Cre+) mice displayed a substantive increase in lung injury that significantly reduced survival during *E. coli*-induced pneumonia, a Gram-negative bacterium that causes severe injury though bacteria are rapidly cleared (already ∼50% clearance by 6H) [37]. The protective effects of neutrophil LOX-1 were also measured in response to *S. pneumoniae*, where LOX-1 limits neutrophil recruitment, potentially via regulation of C3aR and CXCR2, or their cognate chemoattractants, C3a, CXCL1, and CXCL2. Taken together, the current study elaborates on previously identified differences in LOX-1+ neutrophils [22] and solidifies the role of LOX-1 on neutrophils as primarily anti-inflammatory, but not immunosuppressive.

Neutrophils are the first line of recruited defense against pathogens, historically considered a homogenous cell type: short-lived, highly antimicrobial, and potentially tissue-damaging, with little heterogeneity or nuance in their response. However, research advances such as single cell sequencing have expanded our understanding of neutrophils, including subsets with diverse functions, including, but not limited to antigen presentation [8, 38, 39], tissue repair [15, 40, 41], and innate memory [42, 43]. Markers that define these heterogenous populations are continuously being identified, possibly including LOX-1 [22, 44]. While LOX-1 is most recognized for its injurious effects in the context of vascular inflammation [23, 45], we discovered the pulmonary airspaces as a unique niche wherein LOX-1 is essential for limiting lung injury, with emigrated neutrophils as a likely major source [22]. This seemingly contradictory function of LOX-1 as a tissue-protective factor coincides with more recent evidence of its function in the tumor microenvironment, where LOX-1 has been revealed as a marker for polymorphonuclear myeloid derived suppressor cells (PMN-MDSCs) [23]. PMN-MDSCs limit anti-tumor immunity and promote tumor growth [23, 46, 47], supporting dampening effects of LOX-1 on neutrophils based on tissue-specific context. Indeed, the relationship between LOX-1 and MDSCs has been extended to respiratory infection in humans, based on recent evidence that circulating LOX-1+ MDSCs are elevated in patients with severe COVID-19 [24, 44]. Our current and prior [22] investigations reveal that pneumonic airspaces promote neutrophil LOX-1 expression upon transmigration, possibly acting as a negative feedback system to tame activity and limit injury.

Given the contextual role LOX-1+ neutrophils appears to have in disease progression, it is imperative to improve our understanding of how LOX-1 is regulated on neutrophils within the tissue environment. Our current (**Fig 5B**) and published data [22] consistently reveal neutrophils as a prominent source of lung LOX-1 during pneumonia, and a goal of our current study was to validate this finding in humans. We found consistency between human and mouse data, where over half of neutrophils in pneumonic lungs were LOX-1+ (**Fig 1D**), with modest induction between control and pneumonic patients (**Fig 1C**) likely due to the greater degree of overall neutrophilic infiltrates in the latter (**Fig 1B**). Interestingly, our results indicate that lung but not blood neutrophils express LOX-1 (**Fig 2A-B**), suggesting the airspace environment as an essential trigger for LOX-1 induction upon recruitment. In support of this hypothesis, pneumonic BALF was sufficient to trigger LOX-1 induction on naïve bone marrow neutrophils, which have limited surface LOX-1 expression at baseline (**Fig 2C-D**). These data reveal LOX-1 as a defining feature of the lung neutrophil program, possibly contributing to tissue homeostasis.

Our findings also highlight dramatic phenotypic shifts in LOX-1+ neutrophils that are associated with enriched chemokine receptors (**CCR2, CXCR2, CXCR4**, and **CXCR3**) and macrophage-related surface proteins (**MARCO, CD200R, SiglecF**, and **CD68**) (**Fig 3 and Supplemental Fig 1**). Some of these changes were akin to those in cancer-associated PMN-MDSCs and long-lived pro-resolving neutrophils, such as elevated CXCR4 [9], CD200R [48, 49], and SiglecF [31]. Mechanistically, CD200R suppresses TLR signaling through activation of the p120-RasGAP signaling cascade which is ablated under conditions of cellular stress and inflammatory cytokine signaling [50]. MARCO and CD68 have primarily been associated with macrophage physiology and are phagocytic receptors [28, 51], leading to clearance of pathogens [28, 52-55], apoptotic cells [56], and oxidized lipoproteins [57-59], like LOX-1. To our knowledge, our investigation of LOX1+ neutrophils is the first to reveal MARCO induction on lung neutrophils, though recent reports show that like LOX-1, MARCO is highly expressed on MDSCs [60]. Meanwhile, CD68 expression has been identified on neutrophils, but this finding is currently limited to the setting of inflammatory bowel disease, where the function of neutrophil CD68 is unknown [61].

Elevated SiglecF on LOX1+ neutrophils during pneumococcal pneumonia was consistent with our observations of macrophage-centric factors on this neutrophil population and in their immunosuppressive properties. SiglecF was recently identified as a marker for a subset of mature neutrophils, coinciding with CD101 expression [62], that promote fibrosis in the infarcted heart through activation of collagen production [63] or resolve lung inflammation downstream of 12/15-lipoxygenase,17-HDA, and Resolvin D-3 [15]. Another important change observed in LOX-1+ neutrophils was their elevated degree of apoptosis (**Fig 4**). Apoptotic neutrophils represent a major substrate for alveolar macrophage efferocytosis in the wake of acute pulmonary inflammation [64]. Efferocytosis is a key process for inflammatory resolution in the lungs and is more likely to be fueled by LOX-1+ *vs.* LOX-1-neutrophils given the heightened capacity of the former to undergo apoptosis (**Fig 4**). Interestingly, limited studies have identified a direct role for LOX-1 in enhancing efferocytosis [65, 66]. Like recruited neutrophils, resident alveolar macrophages exhibit strong LOX-1 expression [22], along with a prominent role in lung efferocytosis [67], suggesting that LOX-1 may have important roles for both AMs and their targets in the context of lung tissue resolution, constituting an area of future investigation.

In the lungs, both recruited neutrophils and resident alveolar macrophages exhibit high levels of LOX-1 expression. Given our previous work that showed inflammatory and metabolic changes in both cell populations, along with increased lung injury with LOX-1 neutralization [22], we determined neutrophil-dependent functions of LOX-1 during pneumonia. To do so, we leveraged a novel mutant mouse generated by our collaborator Dr. Sawamura containing loxP sites around exon 6 of the *olr1* (LOX-1) gene (**Fig 5A**). These mice were crossed with others bearing a Cre recombinase transgene under the control of the Mrp8 (S100A8/A9) promoter to target neutrophils and were backcrossed to a C57BL/6 background. Following infection with *E. coli,* BALF concentrations of sLOX-1 (soluble LOX-1) and neutrophil surface LOX-1 were considerably reduced and strongly implicate neutrophils as a key source of LOX-1-dependent expression in the pneumonic lung (**Fig 5B-C**). Neutrophil-specific LOX-1 deletion also reduced survival post-*E. coli* infection concomitantly with increased alveolar edema (**Fig 6A-B**), aligning with our prior observations in pneumonic mice following LOX-1 blockade [22]. Moreover, we measured an increase in recruited CD11b^hi^ monocytes that may contribute to inflammatory injury or macrophage replacement (**Fig 6C-E**). To determine whether our findings extend to other causes of lung infection, we infected knockout mice and Cre-littermante controls with a virulent strain of *S. pneumoniae* (serotype 3). Neutrophil LOX-1 deletion significantly increased neutrophil recruitment (**Fig 6F**) and neutrophil-related cytokines (G-CSF, IL-17, and CXCL1) (**Fig 6G**), with strong trends towards increased alveolar edema and bacteremia (**Fig 6H-1**), consistent with our hypothesis that LOX-1 regulates acute pulmonary inflammation. While it has been reported that LOX-1 can improve bacterial adhesion [68, 69], bacterial clearance was unaffected by LOX-1 deletion (**Fig 6J**), indicating that enhanced neutrophilia is unlikely secondary to changes in pathogen burden. In fact, our data from *E. coli-*infected mice (**Fig 4**) indicate increased ROS in LOX-1+ neutrophils, arguing against defects in anti-bacterial defense.

Finally, deletion of LOX-1 on neutrophils altered their phenotype in ways that may result in increased recruitment and/or inflammatory potential. LOX-1 knockout cells had higher expression of the complement receptor, C3aR (**Fig 6F; Supplemental Fig 3**), which may exaggerate C3a-mediated recruitment, though its role in antibacterial immunity remains controversial [70]. We also detected dynamic changes in CXCR4 and CXCR2 (**Fig 6L**; **Fig 7A**), where lower expression of CXCR4 likely promotes bone marrow egress [71] and increased CXCR2 expression further encourages chemotaxis to the tissue [72]. This result is consistent with elevated extravascular neutrophils during *S. pneumoniae* (**Fig 6F**) and, interestingly, suggests that there are pathogen-specific signals that impact neutrophil biology via LOX-1. Taken together, our results provide evidence for LOX-1 as a regulator of neutrophil biology in the lung during pneumonia, where its expression either directly or indirectly limits inflammation via reduced cellular recruitment and potentially other factors, such as improved clearance of apoptotic cells. Follow-up studies will be performed to interrogate the specific mechanisms that facilitate pathogen-specific recovery from pneumonia through neutrophil LOX-1 expression and its downstream outcomes.

## Author contributions

Conceptualization: FTK and LJQ. Methodology: FTK, LJQ, QL, JM, TS, NCC, TB, KTT, and JPM. Investigation: FTK, QL, ESS, MM, DCG, RSI, NH, FS, SNF, AGH, LSF, MRK, MQ, NCC, CSF, AO, and AT. Visualization: FTK, QL, DCG, NCC, AO, and AT. Project administration: FTK and LJQ. Supervision: FTK, LJQ, and NCC. Writing: FTK and LJQ. Writing (review and editing): FTK, LJQ, QL, ESS, JPM, KTT, NCC, TS, and JM. Funding acquisition: FTK, LJQ, JPM, TB, and NCC.

---

**Supplemental Figure 1:**
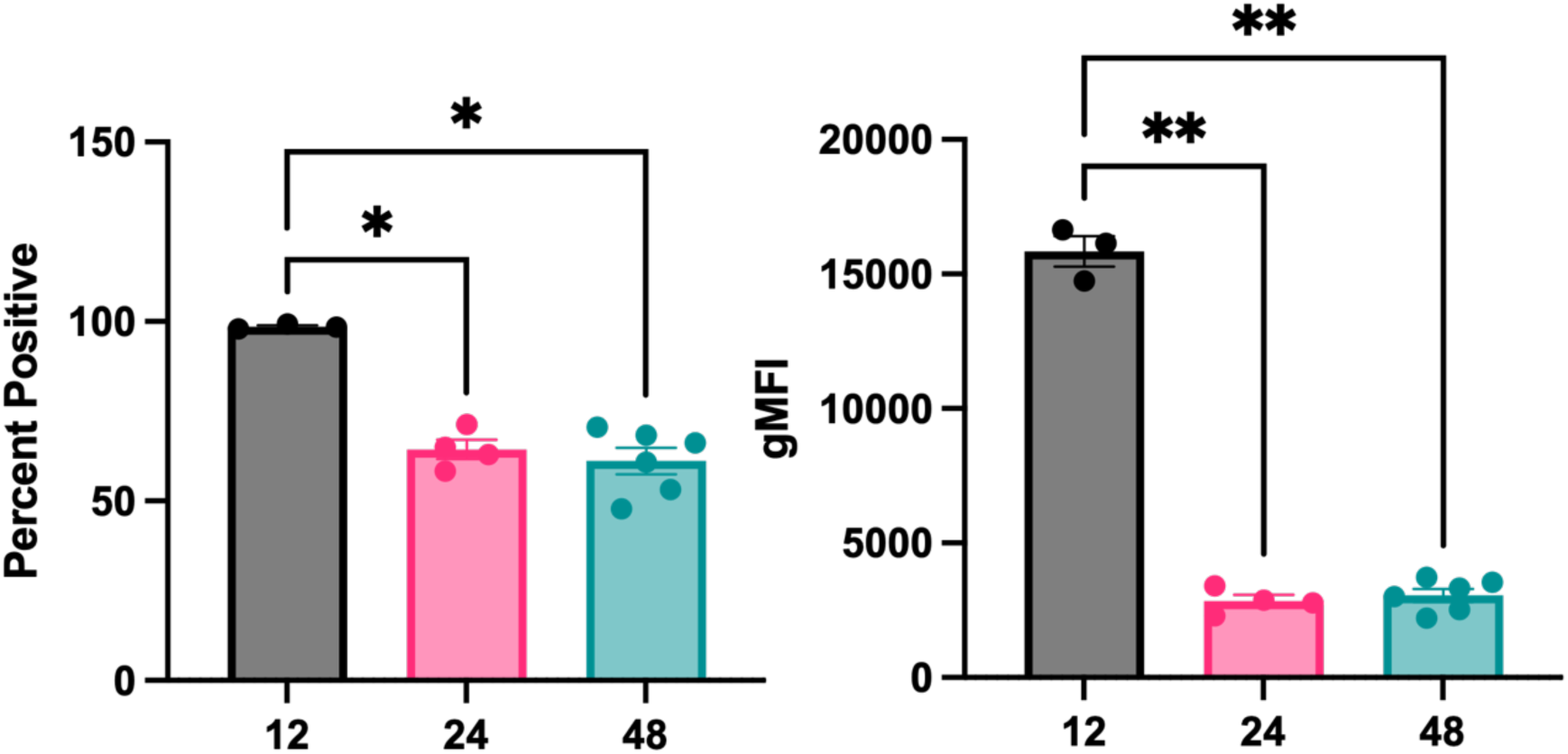
LOX-1+ expression is heterogenous during *S. pneumoniae* infection. Male and female C57BL/6 mice were infected with 1x10^6^ CFU *Streptococcus pneumoniae* (serotype 3) for 12-48H. LOX-1 surface expression was measured by flow cytometry on all neutrophils (Live/CD45+/CD11b+/Ly6G+) isolated from bronchoalveolar lavage fluid (BALF) and fluorescence minus one (FMO) control was used to determine LOX-1 positivity. LOX-1 expression was calculated as percent positive against all neutrophils and level of expression quantified by PE gMFI.

**Supplemental Figure 2:**
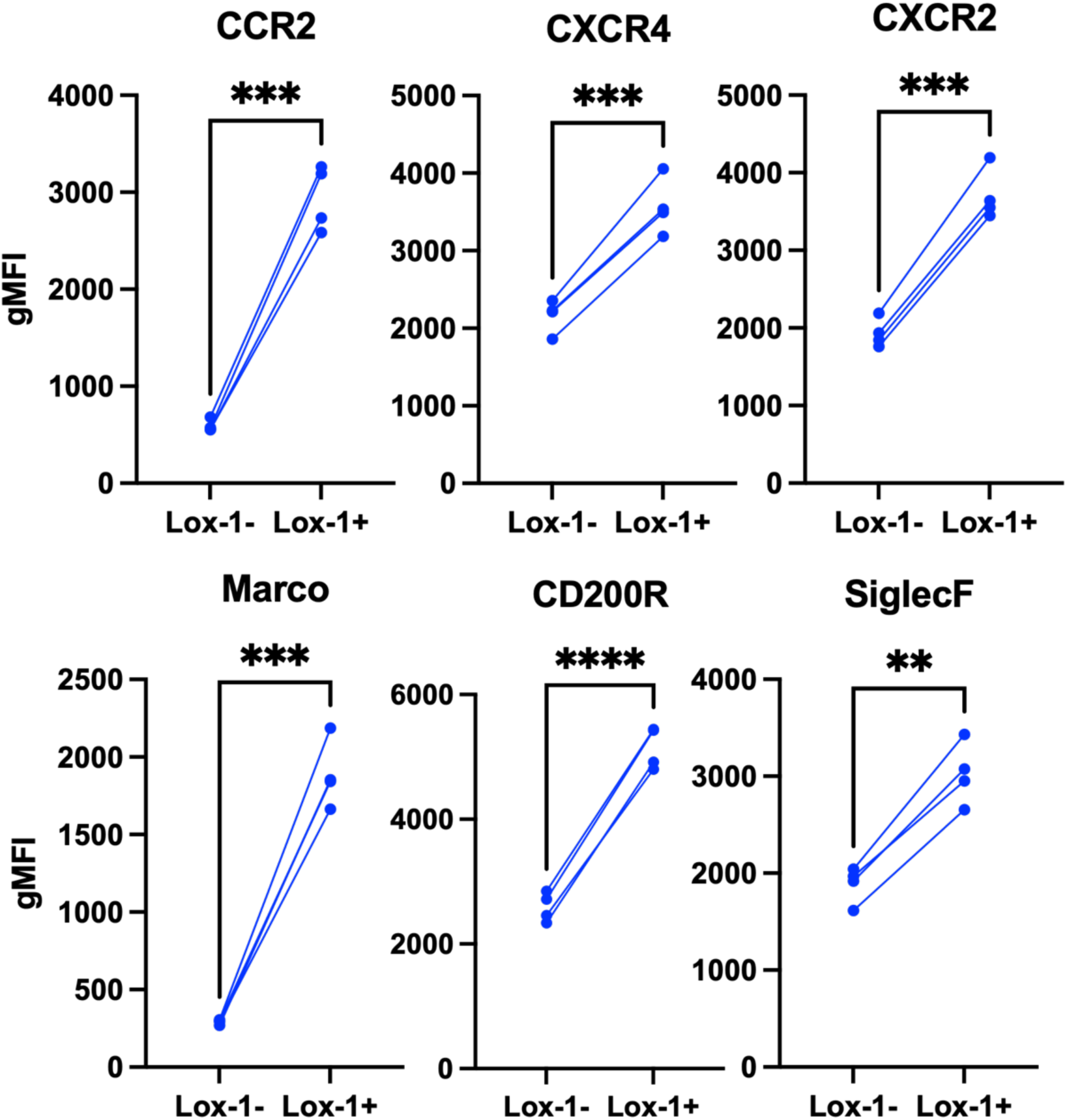
LOX-1+ neutrophils are phenotypically distinct during *S. pneumoniae* infection, consistent with results during *E. coli* infection. Male and female C57BL/6 mice were infected with 1x10^6^ CFU *Streptococcus pneumoniae* (serotype 3). At 24H post-infection chemokine receptors (CCR2, CXCR4, CXCR2), scavenger receptors (CD200R, Marco) and SiglecF surface expression was assessed by flow cytometry on LOX-1+ and LOX-1-neutrophils isolated from BALF using gMFI for quantification. **p<0.01, ***p<0.001, ****p<0.0001 for paired t-test.

**Supplemental Figure 3:**
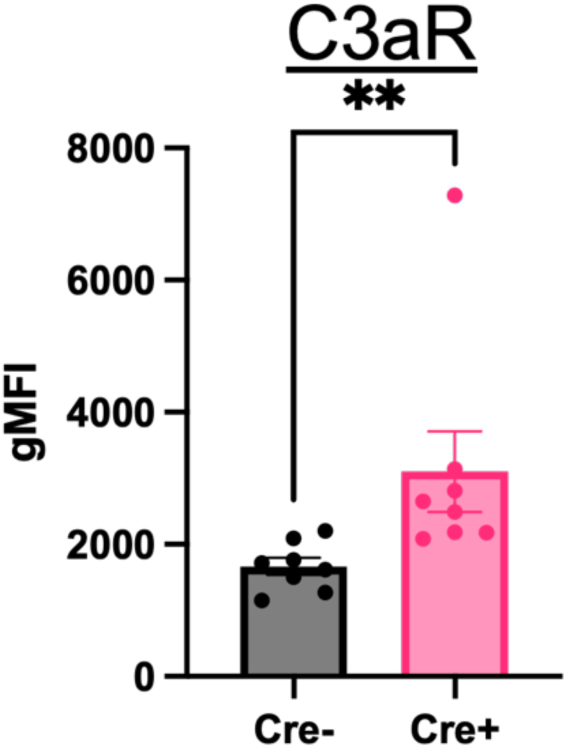
C3aR is elevated in Cre+ neutrophils at 12H post-infection with *E. coli*, akin to *S. pneumoniae* infection. Cre+ and Cre-LOX-1^ΔMRP8^ mice were infected with 1x10^6^ CFU of *E. coli* for 12H. C3aR surface expression was measured on extravascular lung neutrophils (Live/CD45+/Ly6G+/CD11b+) collected from collagenase-digested lung tissue with IV exclusion, described in methods. **p<0.01 for Welch’s t-test.

## References

1. Infections, G.B.D.L.R. and C. Antimicrobial Resistance, Global, regional, and national incidence and mortality burden of non-COVID-19 lower respiratory infections and aetiologies, 1990-2021: a systematic analysis from the Global Burden of Disease Study 2021. Lancet Infect Dis, 2024. 24(9): p. 974-1002.

2. Xiao, K., et al., A pan-immune panorama of bacterial pneumonia revealed by a large-scale single-cell transcriptome atlas. Signal Transduct Target Ther, 2025. 10(1): p. 5.

3. Xie, X., et al., Single-cell transcriptome profiling reveals neutrophil heterogeneity in homeostasis and infection. Nat Immunol, 2020. 21(9): p. 1119–1133.

4. Ravesloot-Chavez, M.M., et al., Tuberculosis susceptibility in genetically diverse mice reveals functional diversity of neutrophils. Elife, 2025. 13.

5. Padhi, A., et al., High-dimensional single-cell analyses reveal neutrophil heterogeneity in guttate psoriasis. EBioMedicine, 2026. 125: p. 106172.

6. Ng, L.G., et al., From complexity to consensus: A roadmap for neutrophil classification. Immunity, 2025. 58(8): p. 1890–1903.

7. Deniset, J.F. and P. Kubes, Neutrophil heterogeneity: Bona fide subsets or polarization states? J Leukoc Biol, 2018. 103(5): p. 829–838.

8. Wu, Y., et al., Neutrophil profiling illuminates anti-tumor antigen-presenting potency. Cell, 2024. 187(6): p. 1422–1439 e24.

9. Qian, J., et al., A CXCR4 partial agonist improves immunotherapy by targeting immunosuppressive neutrophils and cancer-driven granulopoiesis. Cancer Cell, 2025. 43(8): p. 1512–1529 e11.

10. Teijeira, A., et al., CXCR1 and CXCR2 Chemokine Receptor Agonists Produced by Tumors Induce Neutrophil Extracellular Traps that Interfere with Immune Cytotoxicity. Immunity, 2020. 52(5): p. 856–871 e8.

11. Nie, P., et al., A YAP/TAZ-CD54 axis is required for CXCR2-CD44-tumor-specific neutrophils to suppress gastric cancer. Protein Cell, 2023. 14(7): p. 513–531.

12. Salcher, S., et al., High-resolution single-cell atlas reveals diversity and plasticity of tissue-resident neutrophils in non-small cell lung cancer. Cancer Cell, 2022. 40(12): p. 1503–1520 e8.

13. Zhang, Y., et al., Neutrophil subsets and their differential roles in viral respiratory diseases. J Leukoc Biol, 2022. 111(6): p. 1159–1173.

14. Terasaki, A., et al., Neutrophils Expressing Programmed Death-Ligand 1 Play an Indispensable Role in Effective Bacterial Elimination and Resolving Inflammation in Methicillin-Resistant Staphylococcus aureus Infection. Pathogens, 2024. 13(5).

15. Tavares, L.P., et al., Siglec-F(+) neutrophils promote the resolution of acute lung injury through ALOX15 induction. Sci Immunol, 2025. 10(114): p. eaeb2657.

16. Hong, J., et al., gammadelta(+) T-cell-derived IL-17A stimulates airway epithelial/stromal cells to secrete G-CSF, promoting lung-specific pathogenic Siglec-F(+) neutrophil development in PPE-induced emphysema. Cell Mol Immunol, 2025. 22(7): p. 791–805.

17. Hiroki, C.H., et al., Nociceptor neurons suppress alveolar macrophage-induced Siglec-F(+) neutrophil-mediated inflammation to protect against pulmonary fibrosis. Immunity, 2025. 58(8): p. 2054–2068 e6.

18. Podstawka, J., et al., Marginating transitional B cells modulate neutrophils in the lung during inflammation and pneumonia. J Exp Med, 2021. 218(9).

19. Thanabalasuriar, A., et al., PD-L1(+) neutrophils contribute to injury-induced infection susceptibility. Sci Adv, 2021. 7(10).

20. Creusat, F., et al., IFN-gamma primes bone marrow neutrophils to acquire regulatory functions in severe viral respiratory infections. Sci Adv, 2024. 10(41): p. eadn3257.

21. Wang, J.F., et al., Upregulated PD-L1 delays human neutrophil apoptosis and promotes lung injury in an experimental mouse model of sepsis. Blood, 2021. 138(9): p. 806–810.

22. Korkmaz, F.T., et al., Lectin-like oxidized low-density lipoprotein receptor 1 attenuates pneumonia-induced lung injury. JCI Insight, 2022. 7(23).

23. Condamine, T., et al., Lectin-type oxidized LDL receptor-1 distinguishes population of human polymorphonuclear myeloid-derived suppressor cells in cancer patients. Sci Immunol, 2016. 1(2).

24. Hsieh, L.L., et al., SARS-CoV-2 induces neutrophil degranulation and differentiation into myeloid-derived suppressor cells associated with severe COVID-19. Sci Transl Med, 2025. 17(799): p. eadn7527.

25. Shastry, A., et al., Sub-phenotypes of pneumonia defined by pulmonary histopathological features. Am J Respir Crit Care Med, 2026.

26. Mentrup, T., et al., Atherogenic LOX-1 signaling is controlled by SPPL2-mediated intramembrane proteolysis. J Exp Med, 2019. 216(4): p. 807–830.

27. Biocca, S., et al., Simulative and experimental investigation on the cleavage site that generates the soluble human LOX-1. Arch Biochem Biophys, 2013. 540(1-2): p. 9–18.

28. Mukhopadhyay, S., et al., MARCO, an innate activation marker of macrophages, is a class A scavenger receptor for Neisseria meningitidis. Eur J Immunol, 2006. 36(4): p. 940–9.

29. Brunner, J.S., et al., The PI3K pathway preserves metabolic health through MARCO-dependent lipid uptake by adipose tissue macrophages. Nat Metab, 2020. 2(12): p. 1427–1442.

30. Xing, Q., et al., Scavenger receptor MARCO contributes to macrophage phagocytosis and clearance of tumor cells. Exp Cell Res, 2021. 408(2): p. 112862.

31. Pfirschke, C., et al., Tumor-Promoting Ly-6G(+) SiglecF(high) Cells Are Mature and Long-Lived Neutrophils. Cell Rep, 2020. 32(12): p. 108164.

32. Smith, N.M., et al., Regionally compartmentalized resident memory T cells mediate naturally acquired protection against pneumococcal pneumonia. Mucosal Immunol, 2018. 11(1): p. 220–235.

33. Shenoy, A.T., et al., Lung CD4(+) resident memory T cells remodel epithelial responses to accelerate neutrophil recruitment during pneumonia. Mucosal Immunol, 2020. 13(2): p. 334–343.

34. Herjan, T., et al., HuR is required for IL-17-induced Act1-mediated CXCL1 and CXCL5 mRNA stabilization. J Immunol, 2013. 191(2): p. 640–9.

35. Chen, K., et al., IL-17 Receptor Signaling in the Lung Epithelium Is Required for Mucosal Chemokine Gradients and Pulmonary Host Defense against K. pneumoniae. Cell Host Microbe, 2016. 20(5): p. 596–605.

36. Ye, P., et al., Requirement of interleukin 17 receptor signaling for lung CXC chemokine and granulocyte colony-stimulating factor expression, neutrophil recruitment, and host defense. J Exp Med, 2001. 194(4): p. 519–27.

37. Mizgerd, J.P., J.J. Peschon, and C.M. Doerschuk, Roles of tumor necrosis factor receptor signaling during murine Escherichia coli pneumonia. Am J Respir Cell Mol Biol, 2000. 22(1): p. 85–91.

38. Moffat, A. and E. Gwyer Findlay, Evidence for antigen presentation by human neutrophils. Blood, 2024. 143(24): p. 2455–2463.

39. Wang, H., et al., Neutrophil Antigen Presentation Reprograms the Tumor Microenvironment and Elicits Durable and Broad Antitumor Immunity. Cancer Res Commun, 2026. 6(3): p. 500–510.

40. Fischer, A., et al., Neutrophils direct preexisting matrix to initiate repair in damaged tissues. Nat Immunol, 2022. 23(4): p. 518–531.

41. Rizo-Tellez, S.A. and J.G. Filep, Beyond host defense and tissue injury: the emerging role of neutrophils in tissue repair. Am J Physiol Cell Physiol, 2024. 326(3): p. C661–C683.

42. Kalafati, L., et al., The role of neutrophils in trained immunity. Immunol Rev, 2023. 314(1): p. 142–157.

43. Khan, N., et al., beta-Glucan reprograms neutrophils to promote disease tolerance against influenza A virus. Nat Immunol, 2025. 26(2): p. 174–187.

44. Combadiere, B., et al., LOX-1-Expressing Immature Neutrophils Identify Critically-Ill COVID-19 Patients at Risk of Thrombotic Complications. Front Immunol, 2021. 12: p. 752612.

45. Akhmedov, A., et al., Lectin-like oxidized low-density lipoprotein receptor-1 (LOX-1): a crucial driver of atherosclerotic cardiovascular disease. Eur Heart J, 2021. 42(18): p. 1797–1807.

46. Chai, E., L. Zhang, and C. Li, LOX-1+ PMN-MDSC enhances immune suppression which promotes glioblastoma multiforme progression. Cancer Manag Res, 2019. 11: p. 7307–7315.

47. Kim, H.R., et al., The Ratio of Peripheral Regulatory T Cells to Lox-1(+) Polymorphonuclear Myeloid-derived Suppressor Cells Predicts the Early Response to Anti-PD-1 Therapy in Patients with Non-Small Cell Lung Cancer. Am J Respir Crit Care Med, 2019. 199(2): p. 243–246.

48. Choueiry, F., et al., CD200 promotes immunosuppression in the pancreatic tumor microenvironment. J Immunother Cancer, 2020. 8(1).

49. Kim, Y.S., et al., CD200R(high) neutrophils with dysfunctional autophagy establish systemic immunosuppression by increasing regulatory T cells. Cell Mol Immunol, 2024. 21(4): p. 349–361.

50. van der Vlist, M., et al., Signaling by the inhibitory receptor CD200R is rewired by type I interferon. Sci Signal, 2021. 14(704): p. eabb4324.

51. Ng, H.P., et al., Identification of macrosialin (CD68) on the surface of host macrophages as the receptor for the intercellular adhesive molecule (ICAM-L) of. International Journal for Parasitology, 2009. 39(14): p. 1539–1550.

52. Arredouani, M.S., et al., MARCO is the major binding receptor for unopsonized particles and bacteria on human alveolar macrophages. J Immunol, 2005. 175(9): p. 6058–64.

53. Bowdish, D.M., et al., MARCO, TLR2, and CD14 are required for macrophage cytokine responses to mycobacterial trehalose dimycolate and Mycobacterium tuberculosis. PLoS Pathog, 2009. 5(6): p. e1000474.

54. Xu, J., et al., Scavenger Receptor MARCO Orchestrates Early Defenses and Contributes to Fungal Containment during Cryptococcal Infection. J Immunol, 2017. 198(9): p. 3548–3557.

55. Dorrington, M.G., et al., MARCO is required for TLR2- and Nod2-mediated responses to Streptococcus pneumoniae and clearance of pneumococcal colonization in the murine nasopharynx. J Immunol, 2013. 190(1): p. 250–8.

56. Rogers, N.J., et al., A defect in Marco expression contributes to systemic lupus erythematosus development via failure to clear apoptotic cells. J Immunol, 2009. 182(4): p. 1982–90.

57. Zeibig, S., et al., The Scavenger Receptor CD68 Regulates Platelet Mediated Oxidized Low-Density Lipoprotein (oxLDL) Deposition in Atherosclerotic Vessels at an Early Stage of Atherosclerosis in LDLR(-/-)/ApoBec(-/-) Mice. Cell Physiol Biochem, 2019. 52(4): p. 681–695.

58. Ramprasad, M.P., et al., Cell surface expression of mouse macrosialin and human CD68 and their role as macrophage receptors for oxidized low density lipoprotein. Proc Natl Acad Sci U S A, 1996. 93(25): p. 14833–8.

59. Cheng, C., et al., Recognition of lipoproteins by scavenger receptor class A members. J Biol Chem, 2021. 297(2): p. 100948.

60. Liu, S., et al., MARCO expression on myeloid-derived suppressor cells is essential for their differentiation and immunosuppression. Cell Death Discov, 2025. 11(1): p. 337.

61. Amanzada, A., et al., Identification of CD68(+) neutrophil granulocytes in in vitro model of acute inflammation and inflammatory bowel disease. Int J Clin Exp Pathol, 2013. 6(4): p. 561–70.

62. Stephan, J.K., et al., SiglecF-Expressing Neutrophils Differentiate from Mature Neutrophils Locally in the Heart after Myocardial Infarction. Exp Hematol, 2026: p. 105429.

63. Wang, X., et al., Periodontitis-related myocardial fibrosis by expansion of collagen-producing SiglecF+ neutrophils. Eur Heart J, 2025. 46(23): p. 2223–2238.

64. Better, J., et al., Cell type-specific efferocytosis determines functional plasticity of alveolar macrophages. Sci Immunol, 2025. 10(107): p. eadl3852.

65. Murphy, J.E., et al., LOX-1 scavenger receptor mediates calcium-dependent recognition of phosphatidylserine and apoptotic cells. Biochem J, 2006. 393(Pt 1): p. 107–15.

66. Oka, K., et al., Lectin-like oxidized low-density lipoprotein receptor 1 mediates phagocytosis of aged/apoptotic cells in endothelial cells. Proc Natl Acad Sci U S A, 1998. 95(16): p. 9535–40.

67. Pervizaj-Oruqaj, L., et al., The guardians of pulmonary harmony: alveolar macrophages orchestrating the symphony of lung inflammation and tissue homeostasis. Eur Respir Rev, 2024. 33(172).

68. Shimaoka, T., et al., LOX-1 supports adhesion of Gram-positive and Gram-negative bacteria. J Immunol, 2001. 166(8): p. 5108–14.

69. Slavetinsky, J., et al., Wall Teichoic Acid Mediates Staphylococcus aureus Binding to Endothelial Cells via the Scavenger Receptor LOX-1. ACS Infect Dis, 2023. 9(11): p. 2133–2140.

70. Corcoran, J.A. and B.A. Napier, C3aR plays both sides in regulating resistance to bacterial infections. PLoS Pathog, 2022. 18(8): p. e1010657.

71. Eash, K.J., et al., CXCR4 is a key regulator of neutrophil release from the bone marrow under basal and stress granulopoiesis conditions. Blood, 2009. 113(19): p. 4711–9.

72. Eash, K.J., et al., CXCR2 and CXCR4 antagonistically regulate neutrophil trafficking from murine bone marrow. J Clin Invest, 2010. 120(7): p. 2423–31.

