## Supplemental Files for "Neutrophil lectin-like oxidized low-density lipoprotein receptor (LOX-1) limits inflammatory injury during bacterial pneumonia"

### Conflict of interest

The authors have declared that no conflict of interest exists.

### Funding support

- National Institute of Health (NHLBI) (FTK): K99HL159258; R00HL159258.
- National Institute of Health (NHLBI) (LJQ): R01HL165718
- National Institute of Health Shared Instrumentation Grants (NCC): S10OD030269 and S10OD026983.
- National Institute of Health (NIAID) (JPM): R01AI162850

We are especially grateful to the Banner Sun Health Research Institute Brain and Body Donation Program of Sun City, Arizona for the provision of human biological materials.  The Brain and Body Donation Program has been supported by the National Institute of Neurological Disorders and Stroke (U24 NS072026 National Brain and Tissue Resource for Parkinson’s Disease and Related Disorders), the National Institute on Aging (P30 AG019610 and  P30AG072980, Arizona Alzheimer’s Disease Center), the Arizona Department of Health

Services (contract 211002, Arizona Alzheimer’s Research Center), the Arizona Biomedical Research Commission (contracts 4001, 0011, 05-901 and 1001 to the Arizona Parkinson's Disease Consortium) and the Michael J. Fox Foundation for Parkinson’s Research.

**Figure 1**

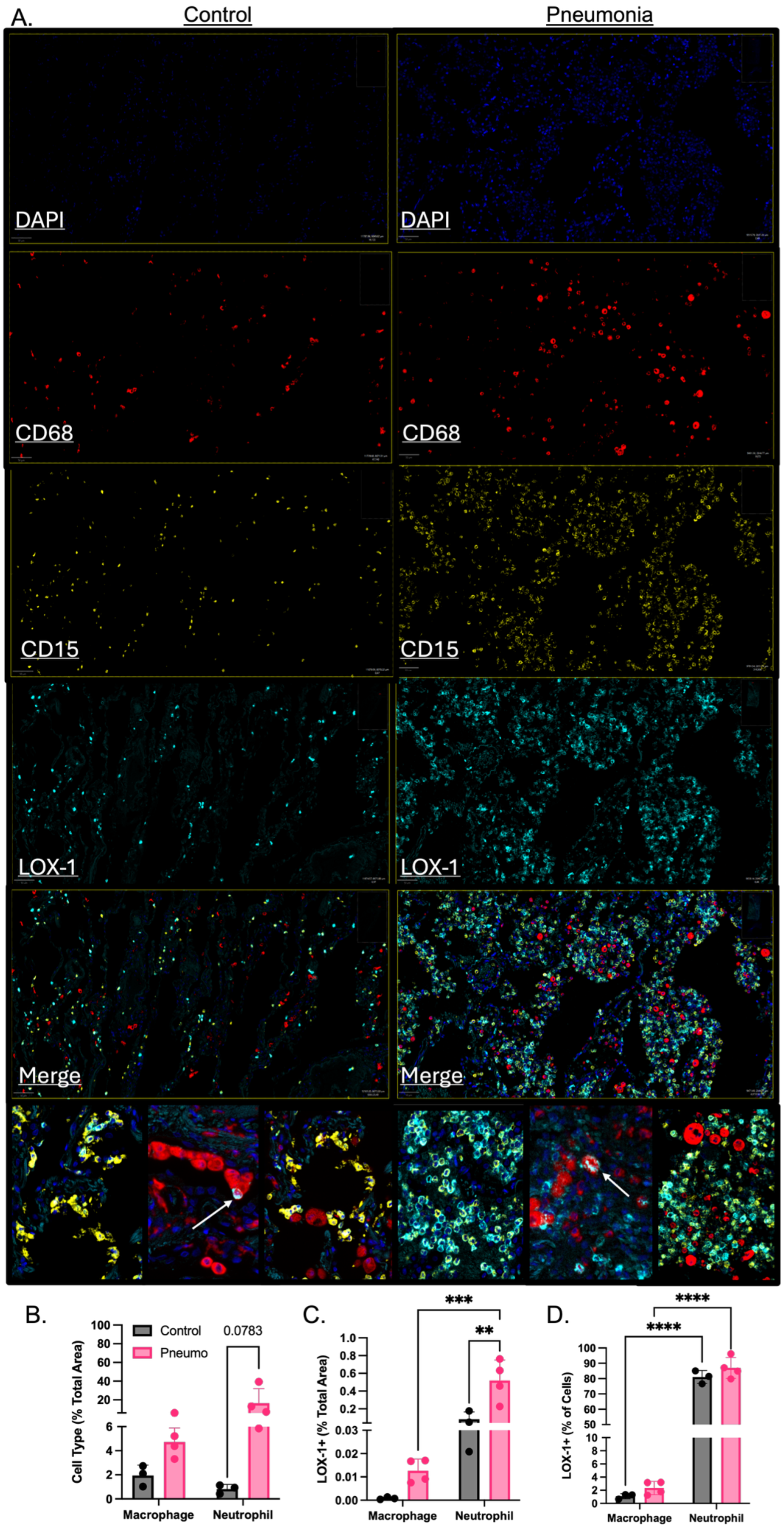
A.
Control
Pneumonia
DAPI
CD68
CD15
LOX-1
Merge
B.
Control
Pneumo
0.0783
Cell Type (% Total Area)
Macrophage
Neutrophil
C.
***
**
LOX-1+ (% Total Area)
Macrophage
Neutrophil
D.
****
****
LOX-1+ (% of Cells)
Macrophage
Neutrophil

**Figure 1: Expression of LOX-1 is detectable on human lung neutrophils regardless of infection status.** A) Rapid autopsies were performed on control or pneumonic human subjects. Immunofluorescence was used to detect LOX-1 (cyan), CD15 (neutrophils; yellow) and CD68 (macrophages; red) on lung sections with DAPI staining nuclei in blue. B) Percent of macrophages and neutrophils were determined against total area (%CD68+ or %CD15+/area analyzed). C) LOX-1 expression was determined on CD68+ cells as % LOX-1+/CD68+ (macrophages) and on CD15+ cells as % LOX-1+/CD15+ (neutrophils) D) LOX-1 positivity by cell type was determined on CD68+ and CD15+ cells as % LOX-1+ out of all CD68+ (macrophage) cells % LOX-1+ out of all CD15+ (neutrophil) cells. Arrows highlight CD68/LOX-1 co-expression. *$p<0.05$ and ***$p<0.001$ for Welch's t-test.

**Figure 2**

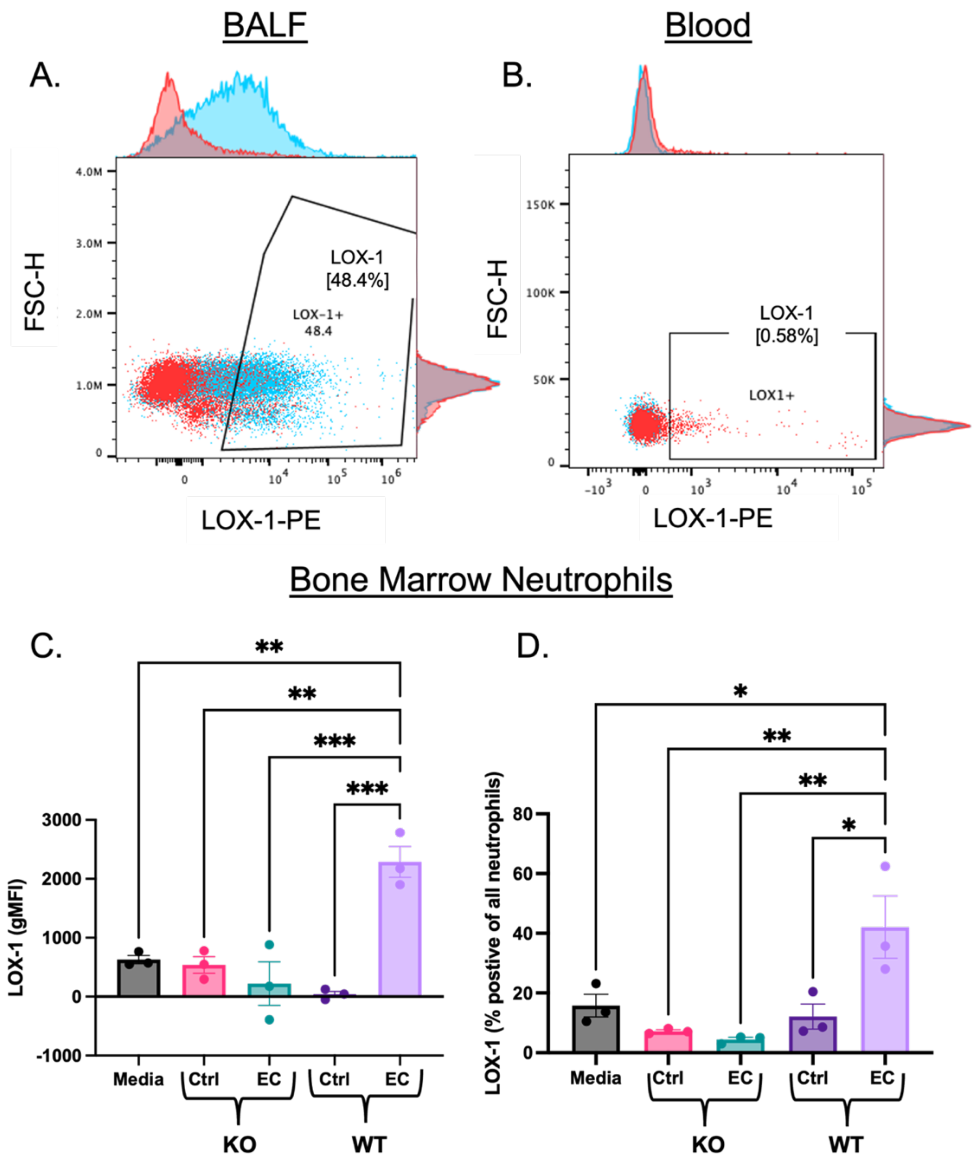

**Figure 2: Neutrophils increase expression of LOX-1 when exposed to the infected airspace.** A) Male and female C57BL/6 mice were infected with 1x10$^{6}$ CFU *Escherichia coli* for 24H. LOX-1 surface expression was measured by flow cytometry on neutrophils (Live/CD45+/CD11b+/Ly6G+) isolated from the A) bronchoalveolar lavage fluid (BALF) and B) heparinized blood. D) Bone marrow neutrophils were isolated from uninfected C57BL/6 and full body LOX-1 knockout (KO) mice and stimulated with media, uninfected BALF or BALF collected from mice infected with *E. coli* for 24H. LOX-1 surface expression was similarly measured at 24H post stimulation *ex vivo*. *p<0.05, **p<0.01, and ***p<0.001 one-way ANOVA with Tukey's multiple comparisons. Representative histograms show LOX-1 expression in sample (blue) and fluorescence minus one (FMO) control (red).

**Figure 3**

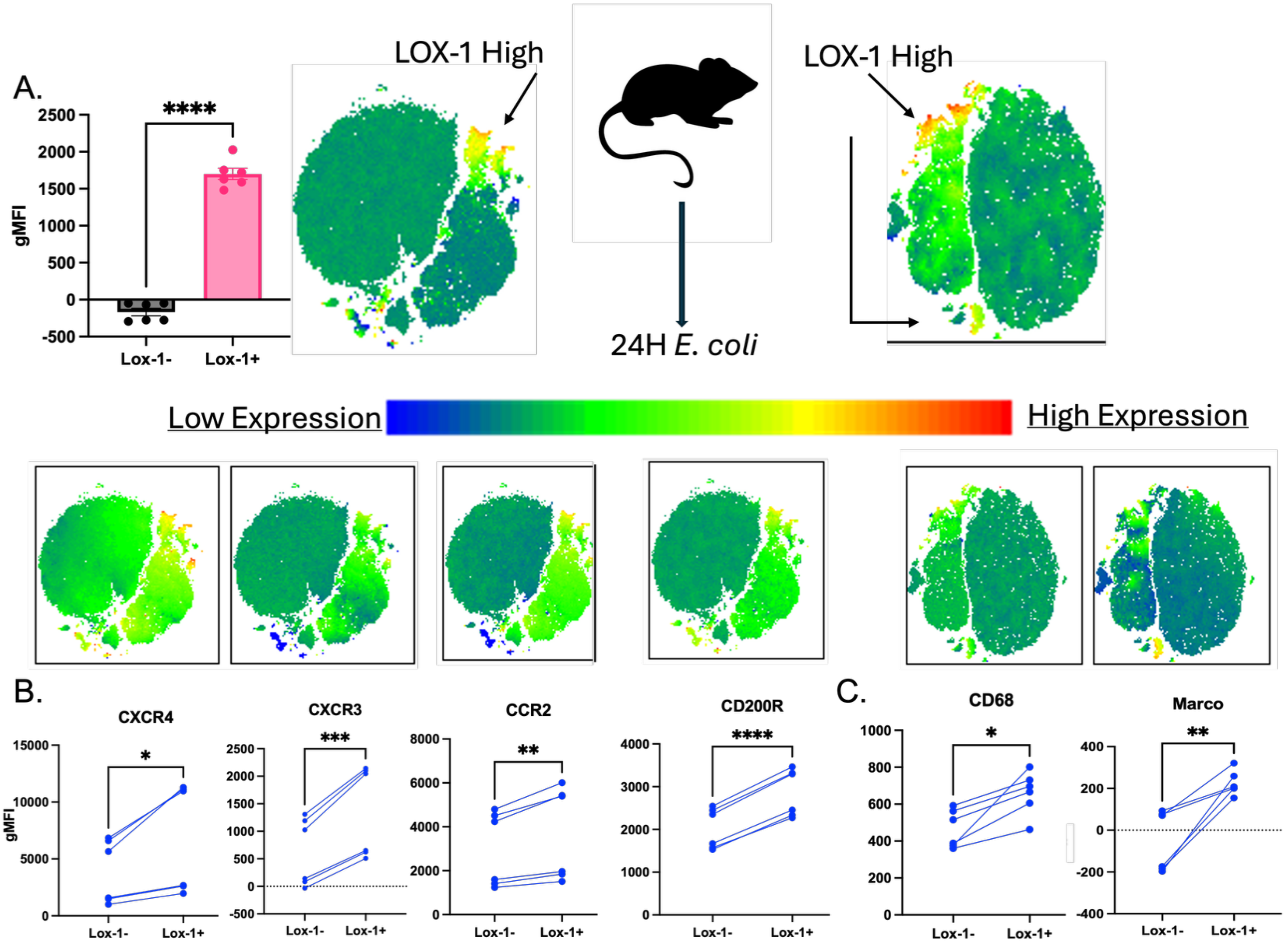

**Figure 3: LOX-1+ neutrophils are phenotypically distinct from LOX-1- cells.** A) Male and female C57BL/6 mice were infected with $1x10^6$ CFU *Escherichia coli* for 24H. LOX-1 surface expression was measured by flow cytometry on neutrophils (Live/CD45+/CD11b+/Ly6G+) isolated from bronchoalveolar lavage fluid (BALF) and fluorescence minus one (FMO) control was used to determine LOX-1 positivity. LOX-1 expression was quantified with PE gMFI and visually shown in yellow-red (high LOX-1) in tSNEs from two separate flow cytometry experiments. B) Chemokine (CXCR4, CXCR3, CCR2) and C) scavenger receptors (CD200R, CD68, Marco) were then assessed by flow cytometry on LOX-1+ and LOX-1- neutrophils using gMFI for quantification and visually represented in tSNEs above each receptor panel. *$p<0.05$, **$p<0.01$, ***$p<0.001$, ****$p<0.0001$ paired t-test.

**Figure 4**

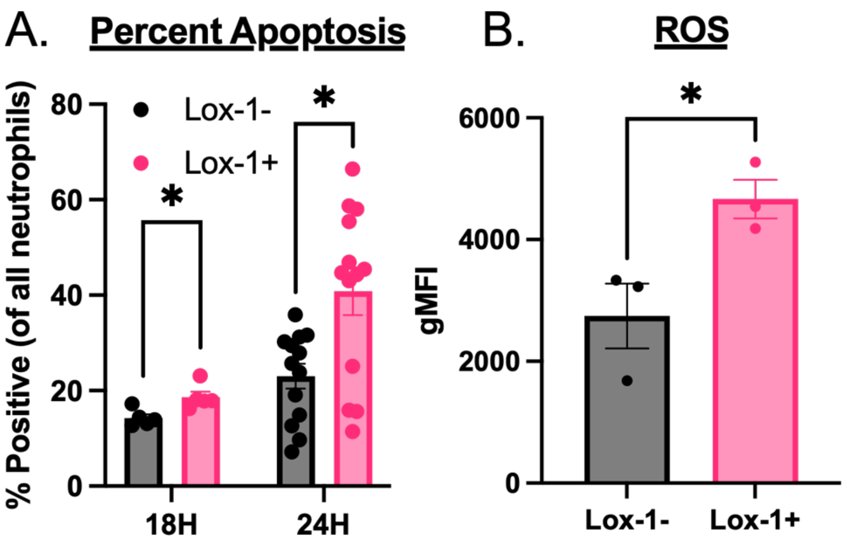

**Figure 4: LOX-1+ neutrophils are more apoptotic and have elevated ROS.** A) Male and female C57BL/6 mice were infected with 1x10$^6$ CFU *Escherichia coli* for 18-24H. LOX-1 surface expression was measured by flow cytometry on neutrophils (Live/CD45+/CD11b+/Ly6G+) isolated from bronchoalveolar lavage fluid (BALF) and fluorescence minus one (FMO) control was used to determine LOX-1 positivity. Early apoptosis was measured with Annexin-5 and propidium iodine (PI) staining where early apoptotic cells are Annexin-5+ and PI. Percent apoptosis was assessed as % of all neutrophils. B) Reactive oxygen species were measured by flow cytometry at 24H post infection with *E. coli.* *$p<0.05$ for paired t-test.

**Figure 5**

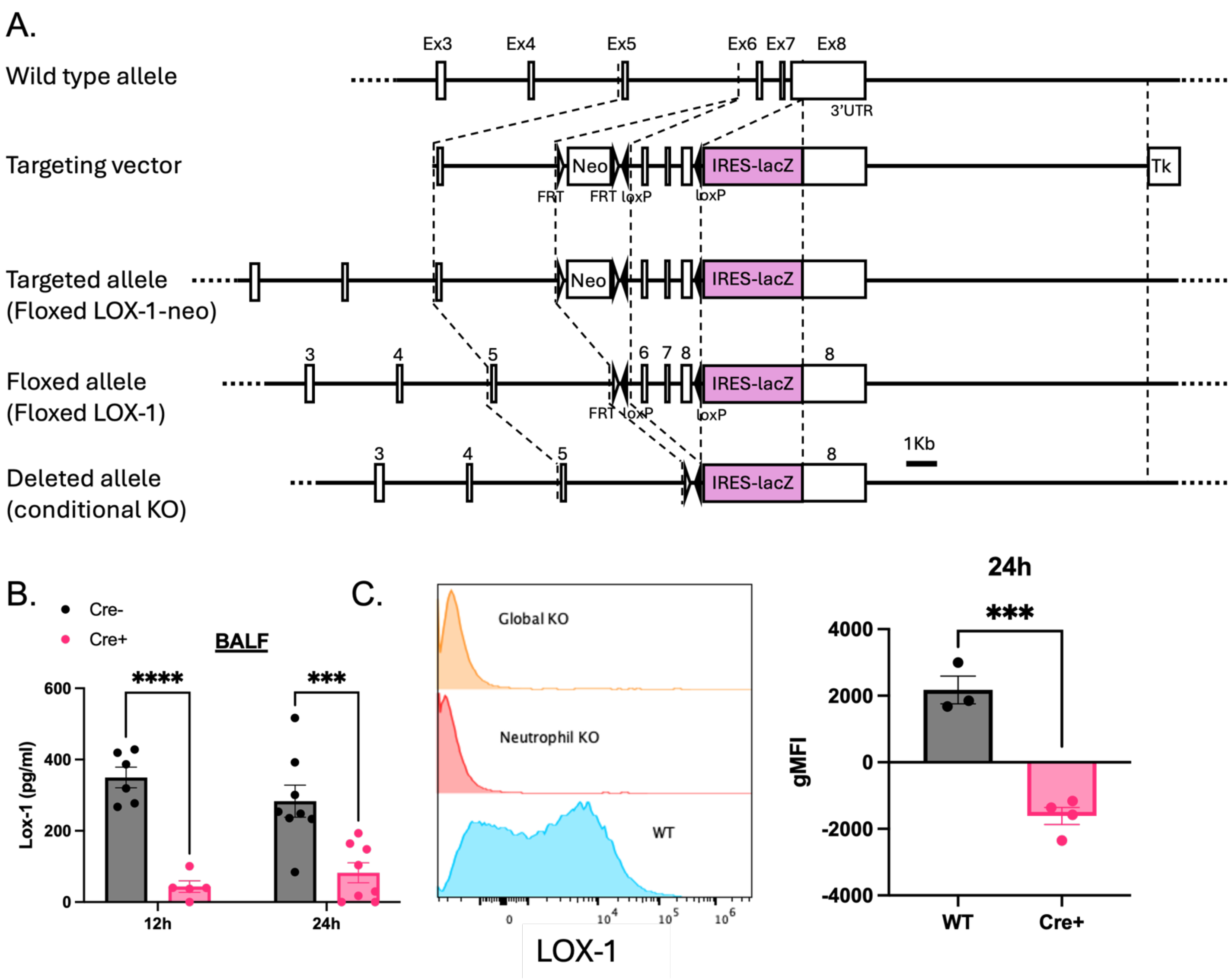

**Figure 5: Generation of conditional neutrophil knockout mouse containing loxP sites surrounding critical exons for LOX-1.** A) A targeting vector was generated to introduce loxP sites around exons 6,7, and an upstream part of exon 8 using the pBluescript phagemid and a LacZ cassette to identify successful clones that were subsequently used to transfect mouse embryonic stem cells. Mice were subsequently crossed with heterozygous Cre-expressing mice under an Mrp8 promoter. B) Cre+ and Cre- LOX-1$^{\Delta MRP8}$ mice were infected with 1x10$^{6}$ CFU of *Escherichia coli* for 12-24H and soluble LOX-1 levels (pg/ml) were determined in BALF at 12 and 24H post-infection. C) LOX-1 surface expression was measured on all neutrophils (Live/CD45+/Ly6G+/CD11b+) at 24H post-infection.

578 **Figure 6**

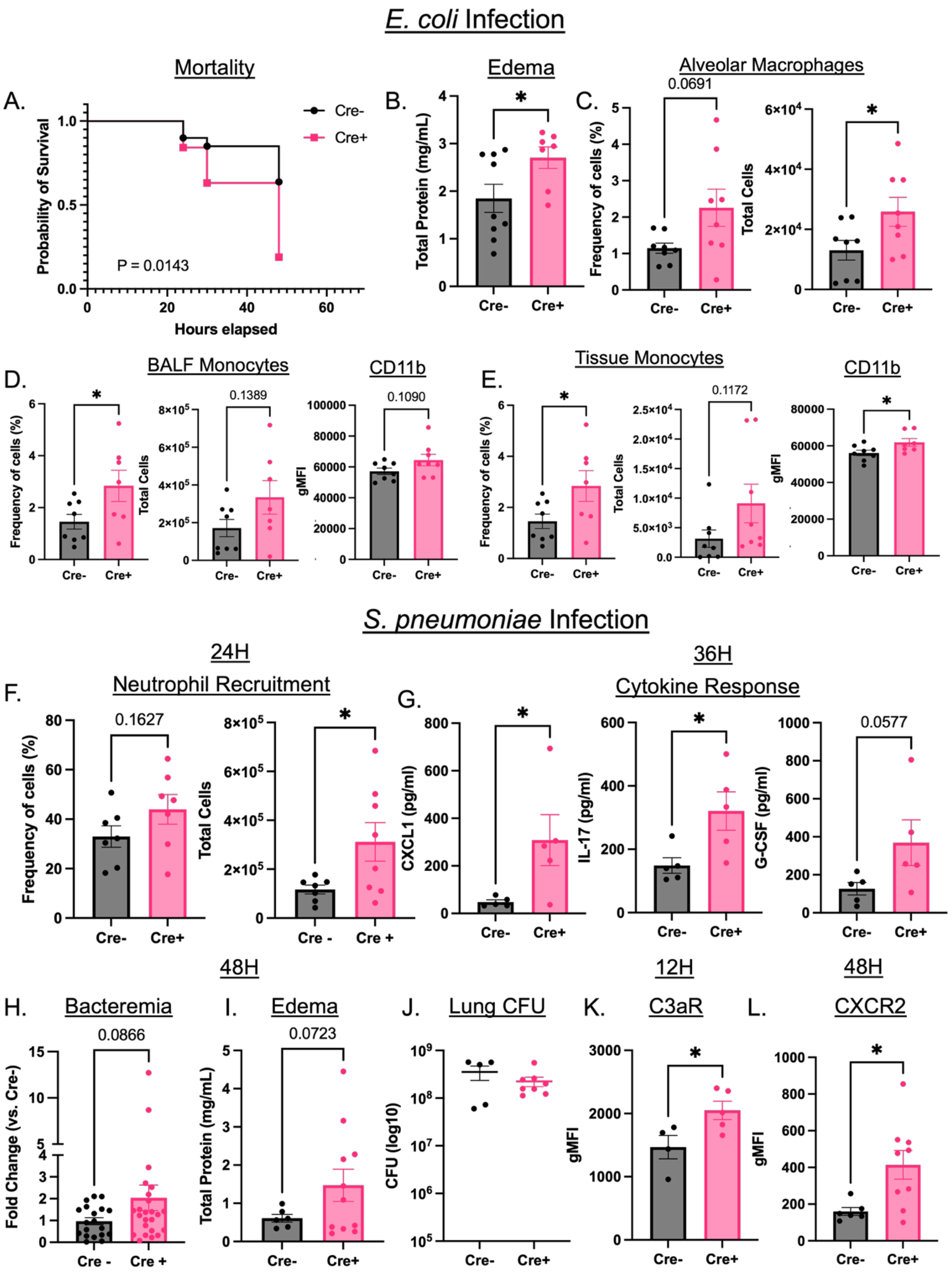

579

**Figure 6: Conditional deletion of LOX-1 in neutrophils exacerbates inflammatory injury and death during pneumonia.** A) Cre+ and Cre- LOX-1$^{\Delta MRP8}$ mice were infected with 1x10$^{6}$ CFU of *Escherichia coli* for 12-48H and survival was assessed up to 48H post-infection. B) total protein (mg/ml) was measured as a surrogate of alveolar edema at 30H post-infection (to limit survival bias). C-E) BALF alveolar macrophages (Live/CD45+/CD11c+/SiglecF+) and extravascular tissue (intravenous anti-CD45.2-) inflammatory monocytes (Live/CD45+/CD11b+/Ly6C+/Ly6G-) frequency and numbers were measured at 12H post-infection along with CD11b expression on D) BALF and E) tissue monocytes. F) Cre+ and Cre- LOX-1$^{\Delta MRP8}$ mice were infected with 1x10$^{6}$ CFU of *S. pneumoniae* (serotype 3) for 12-48H and total BAL neutrophil (Live/CD45+/Ly6G+/CD11b+) numbers and frequency were determined at 24H post-infection along with G) BAL cytokines CXCL1, IL-17, and G-CSF at 36H post-infection. H) Blood bacteria, I) BALF total protein, and J) total lung CFU were determined at 48H post-infection. Frequency of all cells was determined as % of all live cells and backcalculated to the total number of cells. K) C3aR (12H) and CXCR2 (48H) surface expression was measured on extravascular (i.v. CD45.2-) lung neutrophils (Live/CD45+/Ly6G+/CD11b+) collected from collagenase-digested lung tissue. *$p<0.05$, ***$p<0.001$, and ****$p<0.0001$ for student's t-tailed or Welch's t-test. Survival was measured with Log-rank (Mantel Cox) test.

**Figure 7**

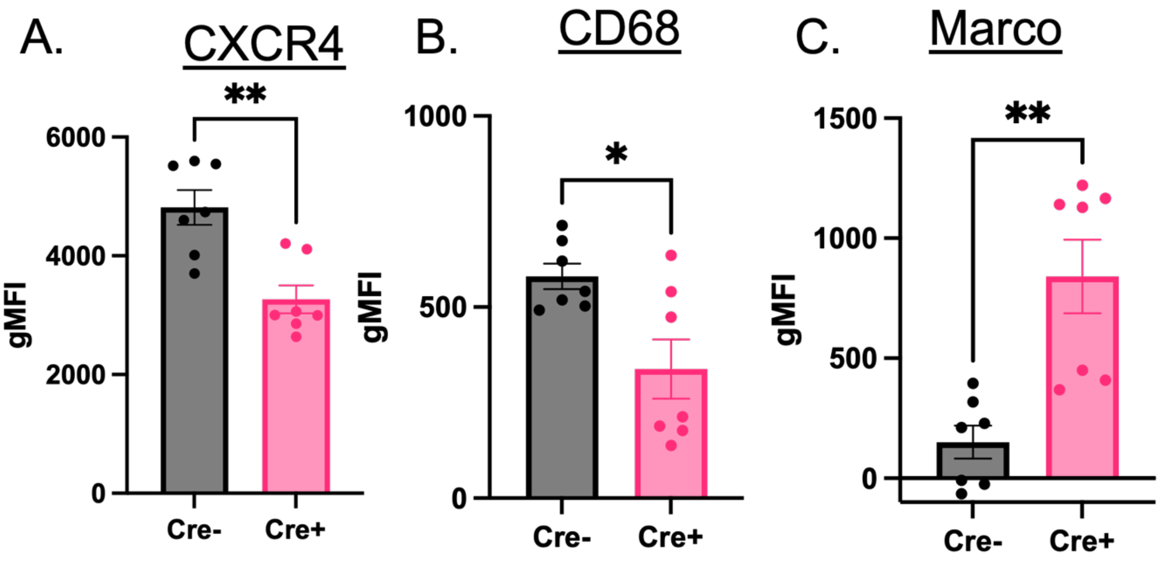

**Figure 7: Conditional deletion of LOX-1 alters neutrophil phenotype during lung infection.** Cre+ and Cre- LOX-1$^{\Delta MRP8}$ mice were infected with 1x10$^{6}$ CFU of *Streptococcus pneumoniae* (serotype 3) for 24H. BALF neutrophils (Live/CD45+/Ly6G+/CD11b+) were isolated and assessed for expression of A) CXCR4, B) CD68, and C) MARCO by flow cytometry. *p<0.05 and **p<01 for student's t-tailed or Welch's t-test.

**Supplemental Figure 1**

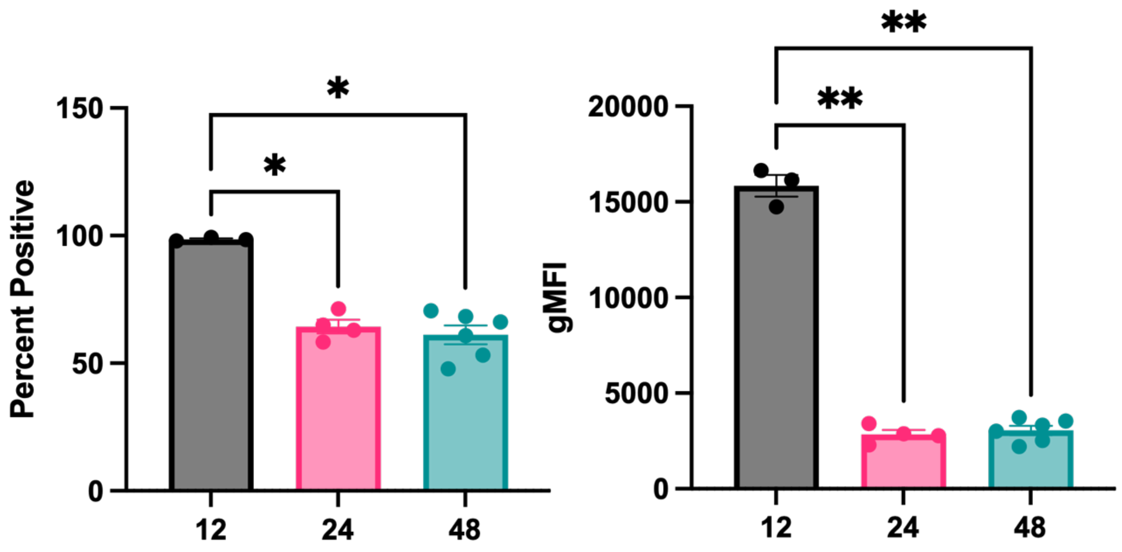

**Supplemental Figure 1: LOX-1+ expression is heterogenous during *S. pneumoniae* infection.** Male and female C57BL/6 mice were infected with $1x10^6$ CFU *Streptococcus pneumoniae* (serotype 3) for 12-48H. LOX-1 surface expression was measured by flow cytometry on all neutrophils (Live/CD45+/CD11b+/Ly6G+) isolated from bronchoalveolar lavage fluid (BALF) and fluorescence minus one (FMO) control was used to determine LOX-1 positivity. LOX-1 expression was calculated as percent positive against all neutrophils and level of expression quantified by PE gMFI.

**Supplemental Figure 2**

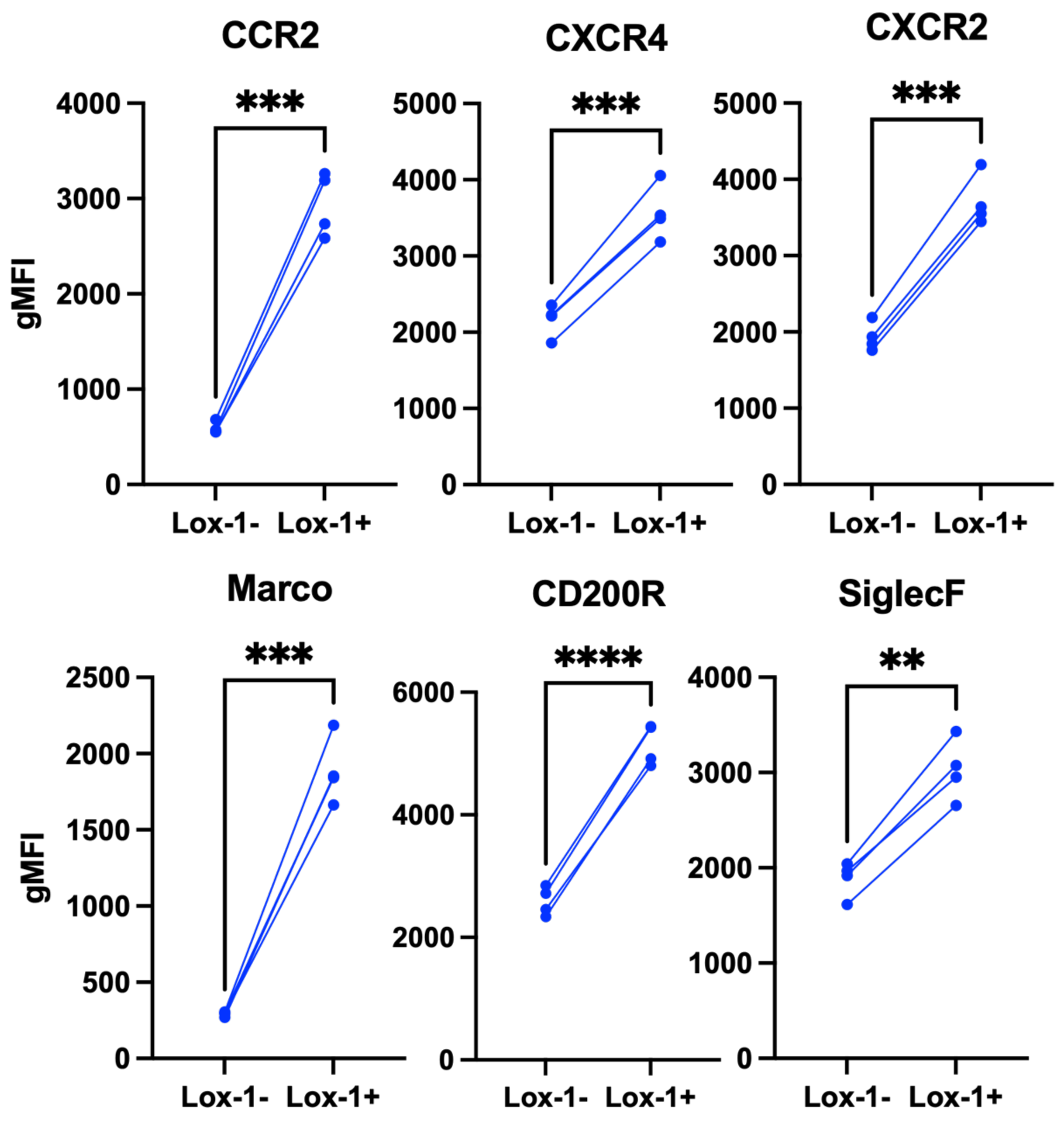

**Supplemental Figure 2: LOX-1+ neutrophils are phenotypically distinct during *S. pneumoniae* infection, consistent with results during *E. coli* infection.** Male and female C57BL/6 mice were infected with $1x10^6$ CFU *Streptococcus pneumoniae* (serotype 3). At 24H post-infection chemokine receptors (CCR2, CXCR4, CXCR2), scavenger receptors (CD200R, Marco) and SiglecF surface expression was assessed by flow cytometry on LOX-1+ and LOX-1- neutrophils isolated from BALF using gMFI for quantification. **p<0.01, ***p<0.001, ****p<0.0001 for paired t-test.

**Supplemental Figure 3**

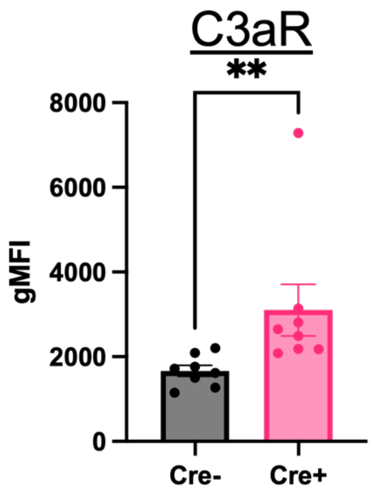

**Supplemental Figure 3: C3aR is elevated in Cre+ neutrophils at 12H post-infection with *E. coli*, akin to *S. pneumoniae* infection.** Cre+ and Cre- LOX-1$^{\Delta MRP8}$ mice were infected with 1x10$^{6}$ CFU of *E. coli* for 12H. C3aR surface expression was measured on extravascular lung neutrophils (Live/CD45+/Ly6G+/CD11b+) collected from collagenase-digested lung tissue with IV exclusion, described in methods. **$p<0.01$ for Welch's t-test.

**Table 1:**

| Sequence | Antigen | Species | Clone | Manufacturer | Catalog | Primary Dilution | Incubation | Fluorophore | Fluorophore Dilution |
|---|---|---|---|---|---|---|---|---|---|
| 1 | Lox1 | Rabbit | E9C5A | CST | 57946 | 1:50 | 37°C, 1 hr | Opal 480 | 1:65 |
| 2 | CD15 | Mouse | HI98 | BD Pharmingen | 555400 | 1:200 | 37°C, 40 min | Opal 570 | 1:150 |
| 3 | CD68 | Rabbit | D4B9C | CST | 76437 | 1:600 | 37°C, 40 min | Opal 690 | 1:100 |

**Table 2**

| Antibody | Clone | Catalog No. |
|---|---|---|
| **Flow Cytometry** | | |
| C3aR-BV480 | 14D4 | BD #753370 |
| CCR2-BUV615 (CD192) | 475301 | BD #751074 |
| CD11b-BUV395 | M1/70 | BD #565976 |
| CD11c-Alexa Fluor 594 | N418 | Biolegend #117346 |
| CD11c-BV785 | N418 | Biolegend #117336 |
| CD200R-Percp-eFluor710 | OX108 | eBioscience #46-5200-82 |
| CD45-Alexa Fluor 594 | 30-F11 | Biolegend #103144 |
| CD45-Alexa Fluor 700 | 30-F11 | Biolegend #103128 |
| CD45-BUV496 | 30-F11 | BD #749889 |
| CD45-BV510 | 30-F11 | Biolegend #103138 |
| CD45.2-BUV737 | 104 | BD #612779 |
| CD68-APC | FA-11 | Biolegend #137008 |
| CXCR2-PE-Cy7 (CD182) | SA044G4 | Biolegend #149316 |
| CXCR3-BV421 (CD183) | CXCR3-173 | Biolegend #126529 |
| CXCR4-BV605 (CD184) | L276F12 | Biolegend #146519 |
| LOX-1-PE | 214012 | R&D #FAB-1564P |
| Ly6C-BUV805 | HK1.4.rMAb | BD #755202 |
| Ly6G-Alexa Fluor 700 | 1A8 | BD #127622 |
| Ly6G-APC-Cy7 | 1A8 | BD #560600 |
| Ly6G-PE-Fire810 | 1A8 | Biolegend #127673 |
| MARCO-Alexa Fluor 594 | 579511 | R&D #FAB2956T |
| MARCO-APC | 579511 | R&D #FAB2956A |
| SiglecF-APC-Cy7 | S17007L | Biolegend #155532 |
| SiglecF-Percp-Fire806 | S17007L | Biolegend #155536 |
| Zombie NIR | | Biolegend #423106 |
| Zombie UV | | Biolegend #423108 |
